# Phylogenetic Dependence and Effective Information in Species-Level Model Evaluation

**DOI:** 10.64898/2026.05.22.727088

**Authors:** Rui Huang, Bin Qi, Deng-Ke Niu

**Author notes:** Correspondence to be sent to: Deng-Ke Niu, College of Life Sciences, Beijing Normal University, Beijing, 100875, China.

## Abstract

Species are widely treated as independent sampling units in comparative analyses, yet shared evolutionary history induces structured dependence that can substantially reduce the amount of independent information available for statistical evaluation and inference. This mismatch between nominal species richness and effective information content can lead to overestimation of statistical precision in species-level analyses. Here, we develop a general framework for quantifying effective information under phylogenetic dependence. We define evaluation subsets embedded within a shared phylogenetic covariance structure and introduce two complementary measures. The first, MIESS (mean-based independence-equivalent sample size), quantifies the amount of independent information available for estimating aggregate quantities under a specified phylogenetic correlation structure, derived from generalized least-squares variance principles. The second, PIESS (prediction-metric-based independence-equivalent sample size), extends this idea to predictive evaluation by mapping uncertainty in standard performance metrics (RMSE, MAE, and *R*²) onto an independence-equivalent sample size scale via calibration against independent-sample benchmarks. We evaluate this framework using empirical mammalian phylogenies (Cricetidae) and idealized tree topologies representing contrasting phylogenetic structures. Across these systems, and under standard models of trait evolution including Brownian motion (BM), Ornstein–Uhlenbeck (OU), and Early-Burst (EB) processes, we compare subsets constructed under phylogenetically dispersed, clustered, and random sampling schemes. Across all settings, dispersed subsets reduce redundancy relative to clustered subsets but do not eliminate dependence-induced information loss. For example, under the *λ*-transformed BM analysis, the dispersed 64-species subset yielded *R*²-based PIESS values well below the nominal size under moderate to strong phylogenetic signal (14.59 at *λ* = 1.00, increasing to 31.96 at λ = 0.50). The resulting loss arises from residual phylogenetic covariance and is robust across evolutionary regimes and sampling fractions. These results indicate that nominal species counts can substantially overestimate independent information in species-level evaluation contexts, and that accounting for phylogenetic dependence is essential for interpreting statistical precision, comparing predictive performance, and designing evaluation protocols in comparative biological analyses.

---

Species-level biological datasets are often described by their nominal number of species, yet shared evolutionary history can make species within an evaluation set internally dependent (Felsenstein 1985). This internal dependence complicates the interpretation of predictive performance in species-level machine learning (Alam et al. 2025, Yu et al. 2025), where evaluation metrics such as root mean squared error (RMSE), mean absolute error (MAE), and *R*²are commonly treated as if derived from independent samples. Phylogenetically informed evaluation designs are increasingly used to reduce training–test leakage by assigning related species to separate blocks and by keeping species closely related to the test set out of the training data (Roberts et al. 2017). For example, in the OGTFinder project, test sets were constructed to exclude genera present in the training set, thereby reducing genus-level overlap between training and evaluation data (Colette et al. 2026). However, such blocking primarily addresses dependence between training and test sets; it does not by itself ensure that species within the evaluation set provide independent evaluation points. Strong internal dependence may therefore remain if the evaluation subset is drawn from a narrow phylogenetic neighborhood.

Such non-independence can substantially reduce the effective information content of an evaluation set (Ané2008, Faes et al. 2009, Bartoszek 2016, Gardner and Organ 2021). When species within a subset are phylogenetically dependent, the nominal species count may considerably overstate the number of effectively independent evaluation points. This loss of effective information can increase the uncertainty of predictive performance evaluation by making performance metrics more sensitive to the particular subset selected. Consequently, predictive performance estimated from species-level evaluation sets may partly reflect similarities arising from shared ancestry rather than independent evidence contributed by distinct evolutionary lineages.

To address these challenges, one practical approach is to select species that are widely separated on the phylogeny, thereby constructing phylogenetically dispersed subsets intended to reduce internal dependence. Building on this idea, we developed a phylogeny-based subsetting framework that adapts subset-selection logic from related fields to the problem of dependence-aware evaluation in species-level machine learning.

The conceptual basis for this subsetting framework draws on prior work in conservation biology, community phylogenetics, and sequence-based studies, where phylogeny-based subset selection has been applied to retain evolutionary history or to choose representative taxa when comprehensive sampling is impractical (Faith 1992, Bordewich et al. 2008, Matsen et al. 2013, Markin et al. 2023). Distance-based metrics such as mean pairwise distance and mean nearest-neighbor distance have been used to characterize phylogenetic clustering or overdispersion (Tucker et al. 2017, Shanker et al. 2026), guiding the construction of dispersed or clustered reference subsets. Here, we adapt this logic to a different purpose: to construct evaluation subsets that can be compared in terms of internal dependence and effective information content.

We present this framework as a dependence-aware evaluation strategy for species-level datasets with phylogenetic structure. The framework combines phylogeny-based subset construction with distance-based metrics, correlation-based diagnostics of internal dependence, and prediction-metric-based independence-equivalent sample size (PIESS). Specifically, we compare dispersed, clustered, and random subsets to evaluate how subset design and alternative covariance assumptions influence within-subset dependence, effective information content, and uncertainty in predictive performance metrics.

## Materials and Methods

### Subset Design for Dependence-Aware Evaluation

We constructed fixed-size evaluation subsets from a phylogenetically defined candidate pool. For a candidate pool *C* containing *N* species, each selected subset was denoted by *S* ⊂ *C*, with target subset size |*S*| = *s*. The same candidate pool was used for all subset types so that distance-based dispersion, clustering, and effective information could be interpreted relative to the phylogenetic structure actually available for sampling.

We compared three subset types. Phylogenetically dispersed subsets were constructed by selecting species from widely separated regions of the phylogeny and were used as favorable low-dependence evaluation subsets. Random subsets of the same size served as empirical baselines representing ordinary sampling from the candidate pool. Phylogenetically clustered subsets were constructed as high-dependence contrast sets, providing reference cases from the opposite end of the dependence gradient.

This design allowed distance-based subset structure and model-implied dependence to be evaluated within the same sampling frame. Specifically, dispersed, clustered, and random subsets were compared under the same *C*, *N*, and *s*, so that differences among subset types reflected subset composition rather than changes in candidate-pool size or sampling fraction.

### Phylogenies and Empirical Candidate Pools

We used both idealized and empirical phylogenetic settings to evaluate how tree topology, candidate-pool size, and subset size influence phylogenetic dispersion, internal dependence, and effective information content in species-level evaluation subsets. As idealized reference cases, we first considered two 32-species ultrametric phylogenies with contrasting topologies: a fully balanced tree and a fully pectinate tree, hereafter referred to as the ladder-like tree. These trees were used to illustrate how subset placement and tree topology influence phylogenetic dispersion and dependence under controlled conditions.

For the empirical analyses, we used Cricetidae as the focal clade. Cricetidae was selected because it provides a moderately large empirical mammalian clade with substantial internal phylogenetic structure, allowing candidate-pool size and subset size to be varied while retaining realistic evolutionary relationships. The clade, containing 528 extant species, was extracted from the DNA-based 4098-species mammalian phylogeny of Upham et al. (2019), accessed through the VertLife mammal tree data portal. From this clade, we randomly selected 512 species to form the largest empirical candidate pool used in the main analysis.

To evaluate the effect of candidate-pool size while preserving a shared empirical phylogenetic context, we constructed a series of nested candidate pools by random down-sampling from the 512-species pool:

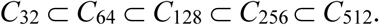

This nested design ensured that smaller candidate pools were contained within larger ones, so that differences among candidate-pool sizes could be interpreted without introducing entirely independent sets of species at each scale.

For each empirical candidate pool, we constructed phylogenetically dispersed subsets and phylogenetically clustered subsets for target subset sizes *s* = 8, 16, 32, and 64, retaining only combinations for which *s* < *N*. For each *N* – *s* combination, we also generated 1000 random subsets of the same size as empirical baselines. Species names and subset memberships for the main empirical subsets are provided in Supplementary Table S1.

### Distance-Based Construction of Dispersed and Clustered Subsets

For each candidate pool *C*, we constructed subsets using patristic distances among species. Let *D* = *d_ij_* denote the patristic distance matrix, where *d_ij_* is the total branch length connecting species *i* and *j*. Because dispersed and clustered subsets represent opposite ends of the phylogenetic-spacing gradient, they were constructed using related but direction-specific distance criteria.

For dispersed subset construction, we summarized within-subset phylogenetic dispersion using three distance-based criteria: the minimum pairwise patristic distance (MinPD), the mean pairwise patristic distance (MeanPD), and the mean nearest-neighbor phylogenetic distance (MeanNND). MinPD captures the distance between the closest pair of species in a subset and is therefore sensitive to local crowding. MeanPD summarizes the overall phylogenetic spread of the subset. MeanNND measures the average distance from each selected species to its nearest selected neighbor and therefore reflects local spacing around selected tips. Together, these criteria describe complementary aspects of dispersion: avoiding extremely close species pairs, maximizing overall spread, and maintaining local separation among selected species.

Phylogenetically dispersed subsets were constructed by prioritizing larger MinPD, with larger MeanPD and larger MeanNND used as secondary and tertiary criteria, respectively. This ordered criterion was chosen because the primary goal of dispersed subset construction was to reduce local phylogenetic redundancy while still favoring subsets broadly distributed across the candidate-pool tree. Subset construction was implemented using a heuristic search procedure consisting of greedy initialization followed by local swap-based refinement, as described in Supplementary Methods Section S1.

Phylogenetically clustered subsets were constructed as high-dependence contrast sets rather than as the primary target of the framework. Because clustered subset construction is inherently tree-local rather than globally dispersive, clustered subsets were not generated by simply reversing the dispersed-subset algorithm. Instead, clustered subsets were constructed using a clustering-oriented distance criterion. Candidate clustered subsets were ranked by smaller MeanPD, with smaller MeanNND and smaller MaxPD used as secondary and tertiary criteria, respectively. Here, MaxPD denotes the maximum pairwise patristic distance within the subset and serves as a boundary-distance criterion, limiting how far apart the most distant selected species can be.

### Random Baselines and Empirical Tail Probabilities

We evaluated the selected subsets relative to random baselines defined separately for each candidate pool *C* and target subset size *s*. For each *N*–*s* combination, we generated 1000 random subsets of size *s*. Each random subset was formed by uniformly sampling *s* distinct species from *C*, while species were allowed to appear in multiple random subsets across repeated draws.

For selected dispersed and clustered subsets, empirical one-sided tail probabilities were calculated as (*b* + 1)/(*B* + 1), where *B* = 1000 is the number of random subsets and *b* is the number of random subsets at least as extreme as the selected subset in the expected direction. The +1 correction avoids reporting zero probabilities when the selected subset is more extreme than all random subsets. For dispersed subsets, larger MinPD, MeanPD, and MeanNND values indicate stronger phylogenetic dispersion; therefore, *b* was defined as the number of random subsets with values greater than or equal to the observed value of the dispersed subset. For clustered subsets, smaller MeanPD, MeanNND, and MaxPD values indicate stronger phylogenetic clustering; therefore, *b* was defined as the number of random subsets with values less than or equal to the observed value of the clustered subset. These empirical tail probabilities quantified the deviation of each selected subset from its corresponding random baseline.

### Correlation-Based Dependence Diagnostics and Mean-Based Information Content

For a subset *S* of size *s*, let *V_S_* denote the phylogenetic covariance matrix for the species in *S* under a specified evolutionary covariance model. This matrix contains the marginal variance of each species on its diagonal and the covariances between species in its off-diagonal entries. To express these covariances on a standardized correlation scale, we converted *V_S_* to the corresponding correlation matrix *R_S_*, in which the diagonal entries are normalized to one and the off-diagonal entries represent pairwise phylogenetic correlations:

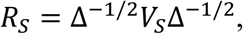

where Δ is the diagonal matrix of marginal variances in *V_S_*. Following the tip-to-tip covariance matrix formulation used in phylogenetic generalized least squares (PGLS, Symonds and Blomberg 2014), this correlation matrix provides a direct summary of tree-induced dependence among the species in the subset.

Brownian motion (BM) was used as the default covariance structure in the main analyses. Under BM, *V_S_* is determined by shared ancestral branch lengths among species. To assess whether the resulting diagnostics were sensitive to covariance assumptions, selected analyses were repeated under *λ*-transformed BM, Ornstein–Uhlenbeck (OU), and early-burst (EB) covariance structures (Pagel 1999, Harmon et al. 2010, Paradis 2014). Pagel’s *λ* transformation rescales the off-diagonal elements of the BM covariance matrix while leaving marginal variances unchanged. OU covariance was parameterized by the phylogenetic half-life ratio *h*/*H*, where *H* is the total tree height. EB covariance was parameterized by a rate parameter *ρ* applied to normalized tree time, *μ* = *t*/*H*, so that branch-specific evolutionary rates were defined as *r*(*μ*) = exp(*ρμ*). Each branch length was replaced by the integral of this rate over the original branch time before the transformed tree was converted to a variance-covariance matrix. All model-implied covariance matrices were then standardized to correlation matrices before calculating dependence diagnostics, including MeanOffCor (mean off-diagonal correlation), MaxOffCor (maximum off-diagonal correlation), MIESS (Mean-based Independence-Equivalent Sample Size), and PIESS. Therefore, alternative models affected the diagnostics through their implied correlation structure rather than through marginal variance scale.

We calculated three phylogenetic-correlation-based diagnostics for each subset. The first is MeanOffCor:

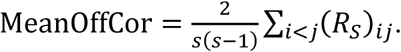

which summarizes the average pairwise correlation induced by shared ancestry within the subset. The second is MaxOffCor:

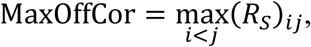

which captures the strongest remaining local dependence. Lower values of both quantities indicate weaker tree-induced dependence. The third diagnostic was MIESS:

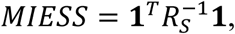

where **1** is a vector of ones. MIESS serves two related purposes in this study. First, it functions as a dependence diagnostic: smaller MIESS values indicate stronger within-subset phylogenetic dependence among species. Second, it translates this dependence into an information-content scale by expressing the subset as the size of an independent sample that would provide the same precision for estimating an overall mean under a generalized least-squares framework with correlation matrix *R_S_* (Aitken 1936, Faes et al. 2009). The derivation and interpretation of MIESS are provided in Supplementary Methods Section S2.

These diagnostics were applied to dispersed, clustered, and random subsets to compare model-implied dependence and mean-based information content across subset types. Trait-level phylogenetic signal metrics such as Pagel’s *λ* and Blomberg’s *K* (Pagel 1999, Blomberg et al. 2003) were not used as within-subset dependence diagnostics because they evaluate how strongly a particular trait conforms to phylogenetic covariance expectations on a given tree rather than quantify the dependence among the species in the subset; this distinction is discussed in Supplementary Methods Section S2.

### Prediction-Metric-Based Independence-Equivalent Sample Size

We performed prediction-metric-based calibration to quantify the effective information available for RMSE, MAE, and predictive *R*²for the selected Cricetidae subsets. Starting from the BM correlation matrix for the 512-species candidate pool, we extracted the rows and columns corresponding to each selected subset, including subsets drawn from nested candidate pools, to obtain the subset-specific correlation matrix used in the simulations. Under the multivariate normal model, this extraction gives the same marginal distribution as simulating the full 512-species tree and retaining only the selected species.

Using the subset-specific BM correlation structure, we simulated true values and prediction errors. True values were generated with marginal variance 1, and prediction errors with marginal variance 0.1, so that prediction performance was neither trivially perfect nor dominated by error variance. Predicted values were defined as the sum of the true values and prediction errors. Each simulation was repeated 10,000 times, and RMSE, MAE, and predictive *R*²were calculated for every replicate. The uncertainty of each metric was summarized as the width of its 95% empirical interval across simulation replicates, consistent with the general expectation that phylogenetic dependence can substantially inflate statistical uncertainty and cannot be ignored in comparative inference (Rabosky 2019). Full details of the simulation procedure are provided in Supplementary Methods Section S3.

To convert this metric uncertainty into an independence-equivalent sample size, we constructed *λ* = 0 benchmarks using independent samples of size *n* = 4, 5, …, 32. These benchmark simulations used the same error variance, number of replicates, and sampling procedure as the phylogenetically structured subset simulations, differing only in the absence of phylogenetic correlation. This matching made the *λ* = 0 simulations suitable benchmarks for translating metric uncertainty under phylogenetic dependence into an independence-equivalent sample size. The PIESS for a subset was then defined as the independent-sample size whose interval width most closely matched that of the subset under the BM correlation structure. This procedure was applied separately to dispersed, clustered, and random subsets.

After PIESS values had been calculated, the random subsets were used as the empirical baseline for evaluating whether the selected dispersed or clustered subset had unusually high or low metric-specific effective information. For dispersed subsets, empirical one-sided tail probabilities were calculated in the direction of higher PIESS than random subsets; for clustered subsets, they were calculated in the direction of lower PIESS than random subsets.

### Sensitivity Analysis under Alternative Covariance Assumptions

Under the primary BM covariance structure (*λ* = 1), we evaluated all empirical *N* – *s* combinations to examine whether the three correlation-based dependence diagnostics and PIESS showed consistent qualitative patterns among dispersed, clustered, and random subsets across candidate-pool and subset sizes. To assess robustness to alternative covariance assumptions, we further evaluated four representative *N* – *s* combinations—*N* = 128, *s* = 8; *N* = 128, *s* = 64; *N* = 512, *s* = 8; and *N* = 512, *s* = 64—under alternative phylogenetic covariance structures, treating these analyses as robustness checks rather than a second full factorial comparison.

For these four representative *N* – *s* combinations, sensitivity analyses were conducted under alternative phylogenetic covariance structures, including *λ*-transformed BM, OU, and tree-height-standardized EB evolutionary models, which are widely used to represent alternative regimes of stabilizing selection, adaptive peak attraction, and time-varying evolutionary rates in phylogenetic comparative analysis (Hansen 1997, Pagel 1999, Butler and King 2004, O’Meara 2012, Paradis 2014). λ-transformed models rescale phylogenetic covariance to vary the strength of phylogenetic signal from complete independence (*λ* = 0) to the original Brownian-motion structure (*λ* = 1), and are therefore used as a standard baseline for assessing dependence strength. OU models represent stabilizing selection toward adaptive optima, with variation in phylogenetic signal determined by the strength of attraction to the optimum (Butler and King 2004). The EB analyses used normalized tree time *μ* = *t*/*H*, making the rate parameter *ρ* dimensionless and comparable across candidate pools with different absolute tree depths. Although early-burst models are not commonly supported in empirical studies (Harmon et al. 2010), we included this regime as an informative boundary case for testing the robustness of the framework to temporally heterogeneous variance accumulation along branches. Analyses were applied to both (i) correlation-based dependence diagnostics, including MIESS, and (ii) PIESS derived from RMSE, MAE, and predictive *R*², quantifying uncertainty in predictive performance. All covariance structures were evaluated after conversion to correlation matrices, and effective-information estimates were interpreted relative to a shared independence benchmark corresponding to *λ* = 0. Full details of covariance parameterizations are provided in Supplementary Methods Section S4.

### Software implementation and reproducibility

All simulations and analyses were performed in R version 4.1.3 (R Core Team 2022) on a Linux system. The proposed phylogenetically informed evaluation workflow was implemented in the PhyloTestESS package version 0.1.0. In this study, the package was used to construct phylogenetically dispersed subsets and clustered reference subsets, generate candidate-pool-constrained random baselines, calculate empirical one-sided *p*-values, compute distance-based subset descriptors, and evaluate covariance-based within-subset dependence diagnostics. The implemented effective-information metrics included MIESS for covariance-based within-subset dependence and PIESS for uncertainty in predictive performance metrics. Details of subset construction and effective-information estimation are provided in Supplementary Methods.

Phylogenetic-tree manipulation and BM covariance calculations were performed using the ape package version 5.8.1 (Paradis and Schliep 2019). Specifically, ape was used to read phylogenetic trees, prune trees to candidate species pools, extract tip numbers, calculate patristic-distance matrices, and construct BM variance-covariance matrices. Principal ape functions used in the workflow included read.nexus(), drop.tip(), Ntip(), cophenetic.phylo(), and vcv.phylo(). Pagel’s *λ* transformation was implemented by rescaling the off-diagonal elements of the BM covariance matrix. OU covariance was calculated from root-to-tip depths, MRCA shared times, and pairwise patristic distances, with the OU parameter expressed as a half-life fraction of total tree height. EB covariance was implemented by first verifying that the tree was rooted and ultrametric, then replacing each branch length by the integral of *r*(*μ*) = exp(*ρμ*) over normalized tree time μ = *t*/*H* before applying vcv.phylo() to the transformed tree. The resulting covariance matrices were standardized to correlation matrices before downstream dependence diagnostics and PIESS simulations.

Standard R packages distributed with R version 4.1.3 were used for routine numerical and graphical operations, including random sampling, seed control, quantile calculation, numerical optimization, matrix-to-correlation conversion, base plotting, and graphical-device control.

## Results

### Illustrative subset configurations on idealized phylogenies

As an illustrative sanity check, we applied the subset-construction procedures to two idealized ultrametric 32-tip phylogenies with clear expected outcomes: a strictly balanced tree and a ladder-like tree. These examples were not intended to represent realistic empirical tree shapes, but to show the qualitative behavior of the dispersed and clustered subset definitions under simple and fully interpretable conditions. On a strictly balanced tree, the dispersed 8-species subset selected one representative from each of the eight major 4-tip clades (Fig. 1a), whereas the clustered reference subset selected species within a single local 8-tip clade (Fig. 1b). In the idealized ladder-like tree, the dispersed subset selected the eight earliest-diverging species along the backbone (Fig. 1c), whereas the clustered reference subset selected the eight most recently diverged terminal species (Fig. 1d). These patterns illustrate that the procedures recover the intended broad-coverage and local-neighborhood configurations under contrasting tree geometries. They also produced the expected differences in information content: under the BM covariance structure with *λ* = 1, MIESS was higher for dispersed subsets than for clustered subsets on both tree geometries (balanced tree: 4.44 vs.1.74; ladder-like tree: 5.70 vs. 1.19).

**Figure 1.**
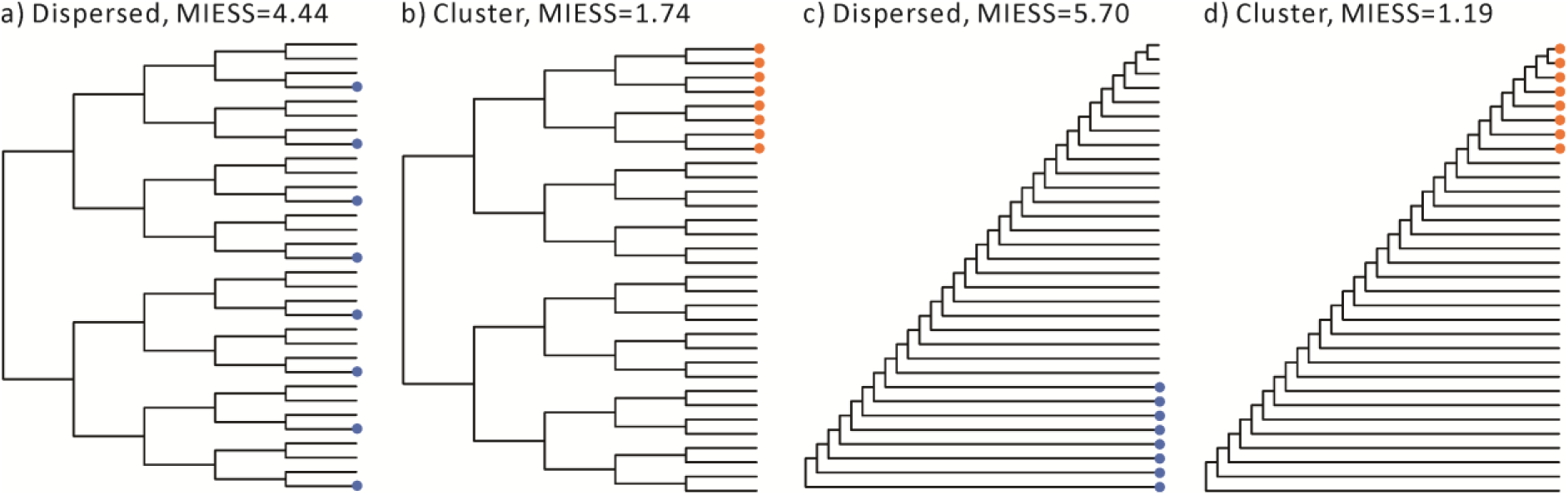
Illustrative subset configurations on idealized phylogenies. Subset-construction procedures were applied to idealized ultrametric 32-tip candidate pools with a target subset size of *s* = 8. The four panels show dispersed and clustered subsets constructed on two contrasting tree geometries: a strictly balanced tree in panels (a) and (b), and a ladder-like tree in panels (c) and (d). Highlighted tips indicate selected species. Values shown in each panel indicate mean-based independence-equivalent sample size (MIESS) calculated from the subset correlation matrix under the Brownian motion covariance structure with *λ* = 1. Graphical layouts are for visualization only; subset construction and evaluation used patristic distances from the underlying trees.

When phylogenetic structure is highly imbalanced or approximates a ladder-like topology, or when early-diverging lineages are sparsely sampled, phylogenetically dispersed subsets tend to converge toward selections of early-diverging taxa along the backbone, making root-proximal sampling a reasonable approximation of maximal phylogenetic coverage in such extreme cases. This behavior is illustrated by the ladder-like tree in Figure 1c, where dispersed sampling reduces to selecting basal lineages along the main evolutionary backbone. Such patterns are most plausibly encountered in systems characterized by strong lineage turnover and serial replacement dynamics, including rapidly evolving viral populations, where RNA virus phylogenies (e.g., influenza A virus and HIV) often exhibit highly unbalanced, ladder-like structures driven by continuous antigenic evolution and selective sweeps (Fitch et al. 1997, Grenfell et al. 2004, Bedford et al. 2012).

### Distance-Based Validation of Dispersed and Clustered Subsets

The empirical validation was conducted on a Cricetidae candidate pool comprising 512 extant species. For this pool, two subsets of equal size (*s* = 64) were constructed: a phylogenetically dispersed subset and a phylogenetically clustered reference subset. In this empirical pool, the selected subsets showed the intended contrast in phylogenetic configuration. The dispersed subset was distributed across widely separated regions of the Cricetidae phylogeny, whereas the clustered reference subset was concentrated within a narrow region of the same phylogeny (Supplementary Fig. S1). Specifically, the clustered reference subset selected 64 of the 66 species in an unnamed local subclade within Arvicolinae. This local subclade showed an approximately ladder-like topology, and the selected species almost completely occupied its most recently diverged portion, closely resembling the behavior observed on the idealized ladder phylogeny in Fig. 1d.

This visual contrast was supported by distance-based comparisons, including MinPD, MeanPD, MeanNND, MaxPD, against 1000 random subsets of the same size (Supplementary Fig. S2). The dispersed subset occupied the upper tails of dispersion-oriented criteria, whereas the clustered reference subset occupied the lower tails of clustering-oriented criteria. Thus, the selected subsets provided suitable low- and high-dependence reference configurations for assessing the consequences of phylogenetic dependence. Detailed metric values, empirical tail probabilities, and distributional comparisons are provided in Supplementary Results Section S1.

### Dispersed Subsetting Reduces Dependence but Retains Limited Effective Information

Having established that the selected subsets showed the intended distance-based contrast relative to random subsets, we next asked whether this contrast also translated into weaker covariance-based dependence and greater effective information in the selected evaluation subsets. For each selected subset and each random subset, we calculated within-subset dependence diagnostics from the subset-specific correlation matrix, *R_S_*, under a BM covariance structure, including MeanOffCor, MaxOffCor, and MIESS. Lower MeanOffCor and MaxOffCor, together with higher MIESS, indicate weaker within-subset phylogenetic dependence.

The phylogenetically dispersed subset showed weaker dependence and higher independence-equivalent information than the random-subset baseline: lower MeanOffCor (0.17 vs. mean = 0.20 and min = 0.16 of random-subsets) and MaxOffCor (0.69 vs. mean = 0.97 and min = 0.89 of random-subsets), and higher MIESS (8.91 vs. mean = 6.79 and max = 8.28 of random-subsets), with one-sided empirical *p*-values of 0.027, 0.001, and 0.001, respectively. The clustered reference subset showed the opposite pattern under the same BM covariance structure (Fig. 2a–c): higher MeanOffCor (0.78 vs. mean = 0.20 and max = 0.31 of random-subsets) and MaxOffCor (0.99 vs. mean = 0.97 and max = 1.00 of random-subsets), and lower MIESS (1.29 vs. mean = 6.79 and min = 5.07 of random-subsets), with *p*-values of 0.001, 0.008, and 0.001, respectively. Thus, the two selected subsets occupied opposite sides of the random-subset distributions not only for patristic-distance metrics, but also for covariance-based summaries of within-subset dependence.

**Figure 2.**
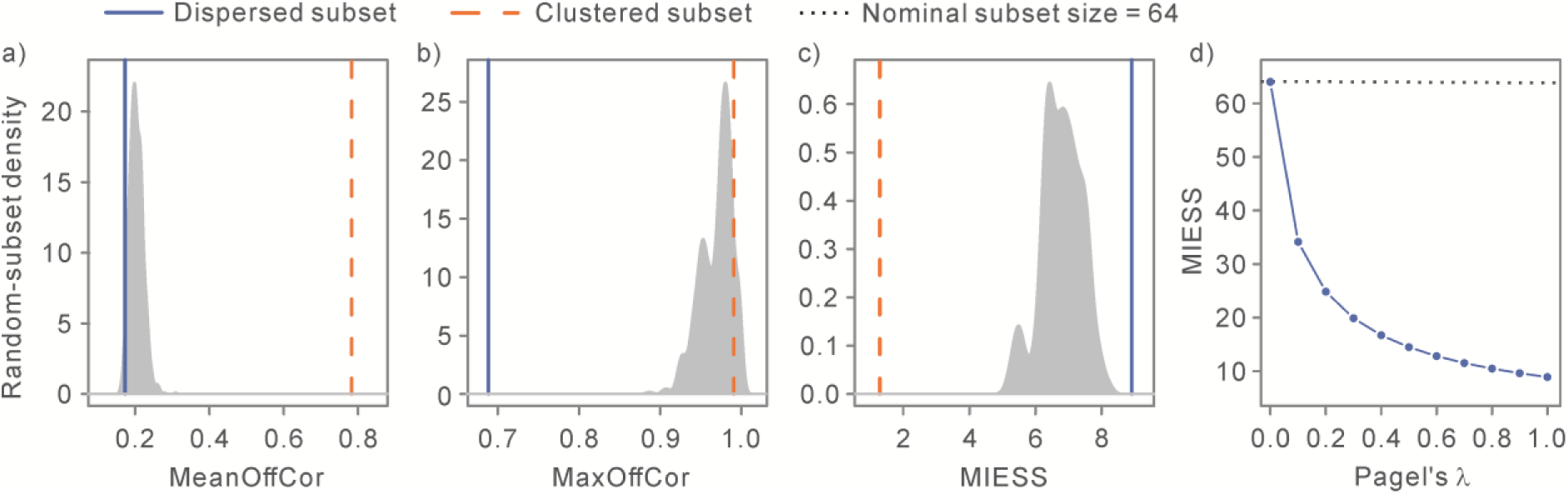
Tree-induced dependence diagnostics for 64-species subsets selected from 512 Cricetidae. Within-subset dependence diagnostics were calculated from the model-implied correlation matrix under a Brownian-motion (BM) covariance structure and compared with empirical random-subset baselines generated from 1000 random subsets. Panels show (a) mean off-diagonal correlation (MeanOffCor), (b) maximum off-diagonal correlation (MaxOffCor), and (c) mean-based independence-equivalent sample size (MIESS). For MeanOffCor and MaxOffCor, smaller values indicate weaker tree-induced dependence. For MIESS, larger values indicate reduced tree-induced dependence, corresponding to greater independence-equivalent information content. Vertical lines indicate observed values for the selected dispersed and clustered subsets, and filled areas show the empirical distributions of the random subsets. Panel (d) shows MIESS of the 64-species dispersed subset under fixed λ-transformed BM covariance structures. The horizontal reference line indicates the nominal subset size.

Although the dispersed subset showed weaker within-subset dependence than both the clustered subset and the random-subset baseline, its MIESS under the BM covariance structure (8.91) remained far below the nominal subset size of 64. We therefore recalculated MIESS for the same 64-species dispersed subset under fixed *λ*-transformed covariance structures (Fig. 2d). MIESS decreased progressively as *λ* increased, from 64.00 at *λ* = 0.0 to 34.14 at *λ* = 0.10, reaching 8.91 at *λ* = 1.0, confirming that the estimator recovers the nominal sample size in the absence of phylogenetic dependence (*λ* = 0.0). This limiting case provided an internal consistency check for the calculation. Thus, the effective sample size of the selected dispersed subset depended strongly on the assumed strength of trait-level phylogenetic dependence.

### Dependence Inflates Uncertainty in Performance Estimation

The MIESS analyses quantified the effective information content implied by the within-subset correlation structure. However, in species-level machine-learning applications, the practical consequence of tree-induced dependence manifests primarily as increased uncertainty (i.e., inflated variance) in predictive performance estimates, rather than merely as patterns in the covariance matrix. To formalize this, we computed the PIESS, which quantifies how many phylogenetically independent samples would be required to match the sampling variance of performance metrics observed for our dispersed evaluation subset. By expressing effective information in terms of estimator variance, PIESS directly links phylogenetic non-independence to the reliability of model evaluation.

Across metrics, PIESS for the 64-species dispersed subset indicated that fewer than 15 independent observations underpinned performance estimation, despite a nominal test size of 64 (Fig. 3). PIESS estimates for RMSE and MAE corresponded to 12.96 and 11.54 independent samples, respectively, while the value for *R*²was slightly higher at 14.36. In contrast, the clustered subset yielded PIESS values below 4 for all metrics, with *R*²again showing the least severe reduction. Although metrics differed modestly within each subset design, the reduction in effective sample size relative to the nominal 64 was pervasive. This comparison demonstrates that dispersed subsetting improved effective information content relative to the clustered configuration, yet remained far short of achieving independence-equivalence to the nominal sample size.

**Figure 3.**
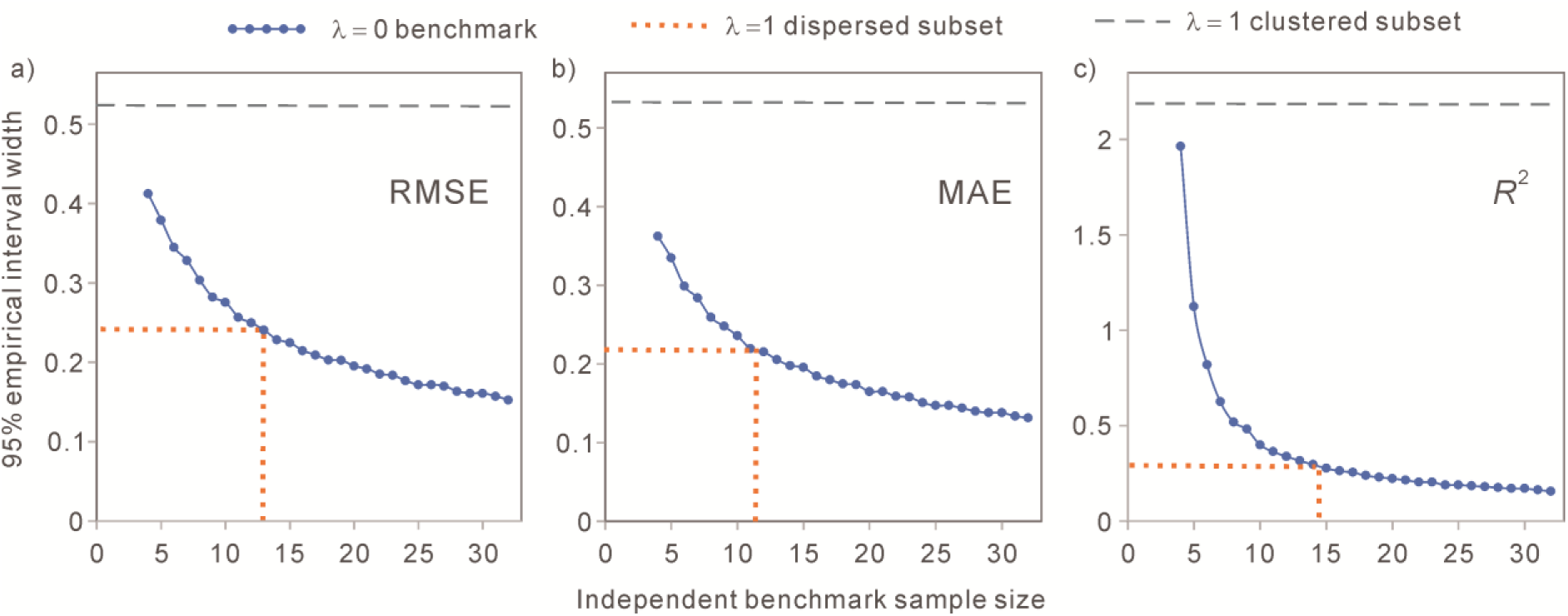
Prediction-metric-based independence-equivalent sample size (PIESS) for dispersed and clustered Cricetidae subsets. Panels (a–c) show the 95% empirical interval width for RMSE, MAE, and *R*², respectively. The solid line, plotted through filled circles corresponding to benchmark sample sizes from 4 to 32, indicates the interval width expected under statistical independence (*λ* = 0). The dotted line and dashed line represent the observed interval widths for the phylogenetic dispersed and clustered subsets, respectively. For the dispersed subset, the intersection with the independence benchmark determines the PIESS (vertical dotted line). Because the clustered subset produced wider intervals than even the smallest benchmark (*n* = 4), its PIESS is recorded as <4.

### Sensitivity of Dependence Across Sampling Designs

We next used the nested Cricetidae candidate pools to evaluate whether the contrast among phylogenetically dispersed, random, and clustered subsets persisted across different candidate-pool sizes and target subset sizes.

Across all feasible *N* – *s* combinations, the distance-based descriptors confirmed that the subset-selection procedure produced the intended phylogenetic contrasts. Phylogenetically dispersed subsets generally showed larger MinPD, MeanPD, and MeanNND than random subsets drawn from the same candidate pool, whereas clustered subsets showed smaller MaxPD, MeanPD, and MeanNND. These patterns were broadly consistent across nested candidate pools and target subset sizes, although the magnitude and statistical strength of the contrasts varied with both absolute subset size and sampling fraction. Notably, the increase in MeanPD for dispersed subsets tended to plateau or become more constrained when the selected subset represented a large fraction of the candidate pool. Detailed distance-based results are provided in Supplementary Results Section S2 and Supplementary Tables S2–S4.

More importantly for the present study, these distance-based contrasts were broadly mirrored by differences in model-implied dependence and independence-equivalent information under the BM covariance structure. Dispersed subsets generally had lower MeanOffCor and MaxOffCor and higher MIESS than random subsets, whereas clustered subsets generally had higher MeanOffCor and lower MIESS (Table 1 and Supplementary Tables S5–S6). These patterns were strongest and most consistent for MIESS, which differed significantly from the random baseline in the expected direction across all analyzed *N* – *s* combinations. MeanOffCor showed the same directional pattern, although some dispersed-subset comparisons were not significant when the selected subset represented a relatively large fraction of the candidate pool. MaxOffCor was less uniformly informative, particularly for clustered subsets, because random subsets could already contain at least one highly correlated species pair. Therefore, the BM-based diagnostics indicate that sampling design affected both within-subset dependence and independence-equivalent information, while also showing that different diagnostics varied in their sensitivity to the dispersed–random–clustered contrast.

**Table 1.** Mean-based independence-equivalent sample size (MIESS) in the nested Cricetidae sensitivity analysis.

| Subset type | $s$ | $N = 32$ | $N = 64$ | $N = 128$ | $N = 256$ | $N = 512$ |
| --- | --- | --- | --- | --- | --- | --- |
| Dispersed | 8 | 5.34*** | 5.60*** | 6.12*** | 6.12*** | 6.20*** |
| Dispersed | 16 | 5.98** | 6.50*** | 7.13*** | 7.19*** | 7.78*** |
| Dispersed | 32 | — | 6.82*** | 7.70*** | 7.80*** | 8.49*** |
| Dispersed | 64 | — | — | 7.85*** | 8.03*** | 8.91*** |
| Clustered | 8 | 1.41*** | 1.19*** | 1.10*** | 1.06*** | 1.05*** |
| Clustered | 16 | 2.06*** | 1.47*** | 1.27*** | 1.13*** | 1.09*** |
| Clustered | 32 | — | 1.92*** | 1.50*** | 1.27*** | 1.14*** |
| Clustered | 64 | — | — | 1.96*** | 1.53*** | 1.29*** |

Values are shown as observed MIESS values. Asterisks indicate empirical one-sided significance relative to random subsets generated from the same candidate pool and with the same target subset size: \* $p < 0.05$ , \*\* $p < 0.01$ , and \*\*\* $p < 0.001$ . For dispersed subsets, significance was evaluated in the direction of higher MIESS than random subsets; for clustered subsets, significance was evaluated in the direction of lower MIESS than random subsets. En dashes indicate combinations not analyzed because subset size $s \geq N$ . The identical dispersed MIESS values for $N = 128$ and $N = 256$ at $s = 8$ reflect recovery of the same selected subset from the nested candidate pools.

The MIESS results are especially relevant to the interpretation of machine-learning evaluation subsets. Although phylogenetic dispersion increased independence-equivalent information relative to random and clustered subsets, the effective information content of dispersed subsets remained far below their nominal species count across the nested Cricetidae analyses. Under the BM covariance structure, MIESS varied systematically with both candidate-pool size and target subset size (Table 1). For dispersed subsets, MIESS generally increased when the target subset size was held constant and the candidate pool became larger, reflecting the greater opportunity to select species from widely separated parts of the phylogeny. When candidate-pool size was held constant, MIESS also increased with target subset size, although the increase was much smaller than the corresponding increase in nominal species count. In clustered subsets, by contrast, MIESS generally decreased with increasing candidate-pool size when target subset size was held constant, because larger candidate pools provided more opportunity to select tightly clustered species. When candidate-pool size was held constant, clustered-subset MIESS increased with target subset size, but remained close to the lower bound expected for strongly dependent samples. Across all configurations, the lowest observed value occurred at *N* = 512 and *s* = 8, where MIESS reached 1.05, illustrating the extreme reduction in effective information under strongly clustered sampling.

Even under one of the most favorable settings for phylogenetic dispersion, where only 8 species were selected from a 512-species candidate pool, the dispersed subset had a MIESS of 6.20, corresponding to 77.5% of its nominal subset size. Thus, strong phylogenetic spacing improved independence-equivalent information, but it did not make the evaluation subset fully equivalent to independent test cases.

### Sensitivity of PIESS Across Sampling Designs

Across all nested Cricetidae sampling configurations, PIESS varied systematically with subset design (Table 2; Supplementary Tables S7–S8). Despite variation in absolute PIESS values across candidate-pool sizes and target subset sizes, the same subset-design pattern was consistently observed: dispersed subsets generally had higher PIESS estimates than random and clustered subsets, whereas clustered subsets consistently produced the lowest PIESS values. PIESS values nevertheless remained well below the nominal subset sizes in all configurations. Thus, the prediction-metric-based analysis supported the same qualitative conclusion as the MIESS results: phylogenetic dispersion increased effective information relative to random and clustered sampling, but did not make the evaluation subsets fully equivalent to independent test cases.

**Table 2.** Prediction-metric-based independence-equivalent sample size (PIESS) for predictive *R*²in the nested Cricetidae sensitivity analysis.

| Subset type | $s$ | $N = 32$ | $N = 64$ | $N = 128$ | $N = 256$ | $N = 512$ |
| --- | --- | --- | --- | --- | --- | --- |
| Dispersed | 8 | 6.89** | 7.05*** | 7.35*** | 7.43*** | 7.46*** |
| Dispersed | 16 | 9.65** | 10.34*** | 10.61*** | 11.10*** | 12.43*** |
| Dispersed | 32 | — | 11.74*** | 12.68*** | 13.27*** | 14.80*** |
| Dispersed | 64 | — | — | 12.37** | 13.76*** | 14.36*** |
| Clustered | 8 | <4*** | <4*** | <4** | <4*** | <4*** |
| Clustered | 16 | 5.96*** | 4.07*** | <4*** | <4*** | <4*** |
| Clustered | 32 | — | 6.20*** | 4.52*** | <4*** | <4*** |
| Clustered | 64 | — | — | 6.44*** | 4.47*** | <4*** |

Values are shown as observed PIESS values for predictive $R^2$ . Asterisks indicate empirical one-sided significance relative to random subsets generated from the same candidate pool and with the same target subset size: \* $p < 0.05$ , \*\* $p < 0.01$ , and \*\*\* $p < 0.001$ . For dispersed subsets, significance was evaluated in the direction of higher PIESS than random subsets; for clustered subsets, significance was evaluated in the direction of lower PIESS than random subsets. En dashes indicate combinations not analyzed because subset size $s \geq N$ .

For dispersed subsets, PIESS increased with both candidate-pool size and target subset size, although the rate of increase remained substantially smaller than the corresponding increase in nominal sample size. This pattern was consistent across RMSE, MAE, and predictive *R*², indicating that phylogenetic dispersion improves prediction-metric stability but does not eliminate dependence-induced loss of effective information. Absolute PIESS values varied across metrics, but all three exhibited the same qualitative dependence structure.

When the target subset size was fixed, PIESS generally increased with greater phylogenetic dispersion, but the magnitude of this increase was limited, especially for small subsets. For example, for *R*²at *s* = 8, increasing the candidate pool from *N* = 32 to *N* = 512, a 16-fold expansion, increased PIESS only from 6.89 to 7.46, corresponding to an 8.27% increase. By contrast, increasing the nominal subset size had a much stronger effect on the absolute amount of prediction-metric-based effective information. Within *N* = 512, increasing *s* from 8 to 64 nearly doubled the *R*²-based PIESS, from 7.46 to 14.36. Thus, although phylogenetic dispersion improved PIESS within a fixed subset size, expanding the number of evaluated species produced a larger gain in independence-equivalent information than increasing dispersion alone. A similar but weaker pattern was observed for MIESS (Table 1), suggesting that prediction-metric-based effective information was more strongly constrained by subset size than the mean-based dependence diagnostic.

In clustered subsets, PIESS values were uniformly low and often approached the lower end of the calibration range, particularly under small target subset sizes and low sampling fractions. This pattern reflects the limited resolution of the PIESS calibration procedure under extreme phylogenetic clustering, where prediction-metric variability under independent benchmarks becomes difficult to distinguish among very small effective sample sizes. Thus, clustered-subset PIESS values should be interpreted as conservative lower-end estimates rather than precise distinctions among very small independence-equivalent sample sizes. For practical model evaluation, however, values near this lower range already provide a strong warning of severe information loss.

### Sensitivity of Dependence Under Alternative Covariance Assumptions

Having evaluated the effect of sampling design under the default BM covariance structure, we next asked whether the core dependence patterns were robust to alternative evolutionary covariance assumptions. We repeated the analyses under *λ*-transformed BM, OU, and EB covariance structures for four representative *N* – *s* combinations: *N* = 128, *s* = 8; *N* = *128*, s = *64*; *N* = 512, *s* = 8; and *N* = 512, *s* = 64 (Table 3; Supplementary Tables S9–S10). These combinations were chosen to span small versus large candidate pools and low versus high sampling fractions without expanding the full *N* – *s* grid. Notably, the setting with *N* = 512, *s* = 64 under *λ* = 1 is the same default covariance setting used in the main analysis shown in Figures 2a-2c, and the default BM configuration is also included in the sensitivity analyses reported in Table 1 and Supplementary Tables S5–S6. These instances are not derived from one another but represent repeated evaluations of an identical benchmark setting across different analytical frameworks. This design choice ensures direct comparability across analyses.

**Table 3.**
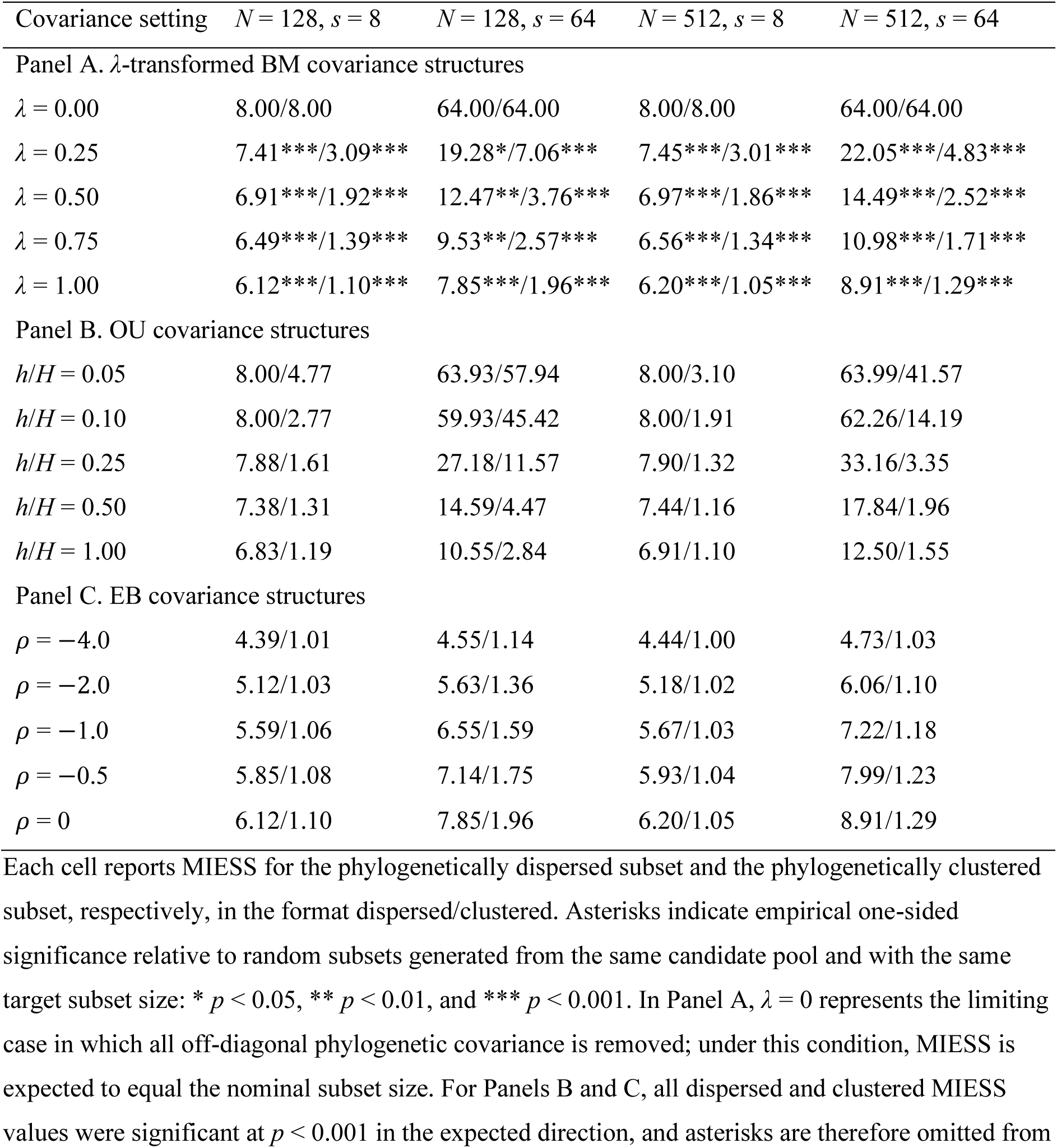
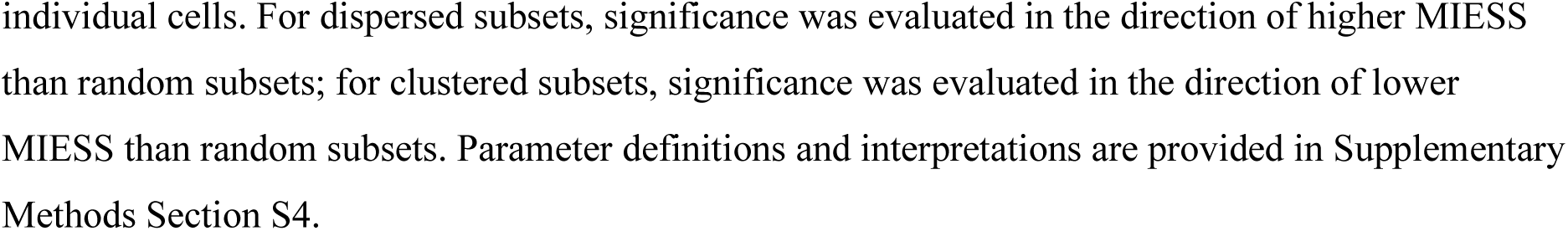
Mean-based independence-equivalent sample size (MIESS) under alternative covariance assumptions in the nested Cricetidae sensitivity analysis.

| Covariance setting | $N = 128, s = 8$ | $N = 128, s = 64$ | $N = 512, s = 8$ | $N = 512, s = 64$ |
| --- | --- | --- | --- | --- |
| Panel A. λ-transformed BM covariance structures | | | | |
| $\lambda = 0.00$ | 8.00/8.00 | 64.00/64.00 | 8.00/8.00 | 64.00/64.00 |
| $\lambda = 0.25$ | 7.41***/3.09*** | 19.28*/7.06*** | 7.45***/3.01*** | 22.05***/4.83*** |
| $\lambda = 0.50$ | 6.91***/1.92*** | 12.47**/3.76*** | 6.97***/1.86*** | 14.49***/2.52*** |
| $\lambda = 0.75$ | 6.49***/1.39*** | 9.53**/2.57*** | 6.56***/1.34*** | 10.98***/1.71*** |
| $\lambda = 1.00$ | 6.12***/1.10*** | 7.85***/1.96*** | 6.20***/1.05*** | 8.91***/1.29*** |
| Panel B. OU covariance structures |  |  |  |  |
| $h/H = 0.05$ | 8.00/4.77 | 63.93/57.94 | 8.00/3.10 | 63.99/41.57 |
| $h/H = 0.10$ | 8.00/2.77 | 59.93/45.42 | 8.00/1.91 | 62.26/14.19 |
| $h/H = 0.25$ | 7.88/1.61 | 27.18/11.57 | 7.90/1.32 | 33.16/3.35 |
| $h/H = 0.50$ | 7.38/1.31 | 14.59/4.47 | 7.44/1.16 | 17.84/1.96 |
| $h/H = 1.00$ | 6.83/1.19 | 10.55/2.84 | 6.91/1.10 | 12.50/1.55 |
| Panel C. EB covariance structures |  |  |  |  |
| $\rho = -4.0$ | 4.39/1.01 | 4.55/1.14 | 4.44/1.00 | 4.73/1.03 |
| $\rho = -2.0$ | 5.12/1.03 | 5.63/1.36 | 5.18/1.02 | 6.06/1.10 |
| $\rho = -1.0$ | 5.59/1.06 | 6.55/1.59 | 5.67/1.03 | 7.22/1.18 |
| $\rho = -0.5$ | 5.85/1.08 | 7.14/1.75 | 5.93/1.04 | 7.99/1.23 |
| $\rho = 0$ | 6.12/1.10 | 7.85/1.96 | 6.20/1.05 | 8.91/1.29 |

Each cell reports MIESS for the phylogenetically dispersed subset and the phylogenetically clustered subset, respectively, in the format dispersed/clustered. Asterisks indicate empirical one-sided significance relative to random subsets generated from the same candidate pool and with the same target subset size: \* $p < 0.05$ , \*\* $p < 0.01$ , and \*\*\* $p < 0.001$ . In Panel A, $\lambda = 0$ represents the limiting case in which all off-diagonal phylogenetic covariance is removed; under this condition, MIESS is expected to equal the nominal subset size. For Panels B and C, all dispersed and clustered MIESS values were significant at $p < 0.001$ in the expected direction, and asterisks are therefore omitted from
individual cells. For dispersed subsets, significance was evaluated in the direction of higher MIESS than random subsets; for clustered subsets, significance was evaluated in the direction of lower MIESS than random subsets. Parameter definitions and interpretations are provided in Supplementary Methods Section S4.

Across covariance models, dependence diagnostics consistently preserved the qualitative ranking induced by sampling design, with phylogenetically dispersed subsets exhibiting lower MeanOffCor and higher MIESS than random subsets, and clustered subsets exhibiting higher MeanOffCor and lower MIESS than random subsets (Table 3 and Supplementary Table S9). MaxOffCor required a more nuanced interpretation (Supplementary Table S10). For dispersed subsets, MaxOffCor consistently showed significantly lower values than random subsets, reflecting the effective removal of highly correlated species pairs under phylogenetic dispersion. For clustered subsets, the pattern depended on subset size. At *s* = 8, clustered subsets exhibited significantly higher MaxOffCor than random subsets, indicating that clustering successfully concentrates strongly correlated species pairs. However, at *s* = 64, clustered subsets were not significantly different from random subsets in terms of MaxOffCor, regardless of candidate-pool size (*N* = 128 or 512). This non-significance does not indicate weak clustering per se, but rather suggests that at larger subset sizes, random subsets already tend to include at least one highly correlated species pair, reducing the discriminative power of the maximum pairwise correlation metric.

Across these representative settings, covariance assumptions strongly affected the absolute magnitude of the dependence diagnostics. Under *λ*-transformed BM, reducing *λ* progressively weakened off-diagonal covariance until, at *λ* = 0, all off-diagonal phylogenetic covariance was removed. This limiting case provided an internal consistency check for the *R_S_*-based diagnostics: as expected, MeanOffCor and MaxOffCor both reached 0.00, whereas MIESS reached the nominal subset size across all evaluated configurations. This confirmed that the diagnostics responded correctly when phylogenetic covariance was eliminated from the model.

The OU and EB analyses showed the same general dependence of the diagnostics on covariance assumptions. Under the OU model, MIESS decreased as the relative half-life increased, indicating stronger retained phylogenetic dependence under larger *h*/*H* values. For instance, for *N* = 512 and *s* = 64, dispersed-subset MIESS declined from 63.99 at *h*/*H* = 0.05 to 12.50 at *h*/*H* = 1.00. Under the tree-height-standardized EB model, negative *ρ* values concentrated variance accumulation toward deeper branches, whereas *ρ* = 0 recovered the BM baseline. After conversion to correlation matrices, this root-weighted variance accumulation generally strengthened shared phylogenetic dependence and reduced MIESS relative to the *ρ* = 0 condition. Therefore, the EB results should be described as a sensitivity analysis of temporal variance weighting along the tree, not as a monotonic relaxation toward independence.

### Sensitivity of PIESS Under Alternative Covariance Assumptions

We further evaluated whether PIESS showed similar robustness to alternative covariance assumptions. Across *λ*-transformed BM, OU, and EB covariance structures, PIESS generally preserved the qualitative ranking induced by subset-construction strategy: phylogenetically dispersed subsets tended to have higher PIESS than random subsets, whereas clustered subsets tended to have lower PIESS than random subsets (Table 4 and Supplementary Tables S11 and S12). This pattern was observed across RMSE, MAE, and predictive *R*², although a small number of comparisons were not statistically significant.

**Table 4.** Prediction-metric-based independence-equivalent sample size (PIESS) for predictive *R*² under alternative covariance assumptions in the nested Cricetidae sensitivity analysis.

| Covariance setting | $N = 128, s = 8$ | $N = 128, s = 64$ | $N = 512, s = 8$ | $N = 512, s = 64$ |
| --- | --- | --- | --- | --- |
| Panel A. λ-transformed BM covariance structures | | | | |
| $\lambda = 0.00$ | 7.74/7.73 | 62.35/62.90 | 7.73/7.90 | 62.48/62.13 |
| $\lambda = 0.25$ | 7.56/6.63*** | 48.82**/34.55*** | 7.85*/6.46*** | 50.81***/24.45*** |
| $\lambda = 0.50$ | 7.56*/4.92*** | 29.84**/17.09*** | 7.80**/4.84*** | 31.86***/10.00*** |
| $\lambda = 0.75$ | 7.59***/<4*** | 18.04**/9.78*** | 7.73***/<4*** | 21.59***/5.92*** |
| $\lambda = 1.00$ | 7.24***/<4*** | 12.84***/6.42*** | 7.37***/<4** | 14.59***/<4*** |
| Panel B. OU covariance structures |  |  |  |  |
| $h/H = 0.05$ | 8.00/6.85*** | 66.12*/62.39 | 7.88/5.85*** | 62.11/56.29 |
| $h/H = 0.10$ | 7.83/5.74*** | 61.24/54.89** | 7.64/4.67*** | 63.17**/37.26** |
| $h/H = 0.25$ | 7.78/<4*** | 56.17***/33.17*** | 7.78/<4*** | 57.49***/11.89*** |
| $h/H = 0.50$ | 7.78**/<4*** | 31.38***/17.17*** | 8.03***/<4*** | 38.61***/6.57*** |
| $h/H = 1.00$ | 7.57**/<4*** | 20.03***/10.03*** | 7.89***/<4*** | 23.62***/4.95*** |
| Panel C. EB covariance structures |  |  |  |  |
| $\rho = -4.0$ | 6.08***/<4 | 5.35/<4*** | 6.33***/<4 | 6.01***/<4*** |
| $\rho = -2.0$ | 6.83***/<4 | 7.30*/4.20*** | 6.81***/<4 | 8.68***/<4*** |
| $\rho = -1.0$ | 7.17***/<4* | 9.24*/4.98*** | 7.09***/<4* | 10.66***/<4*** |
| $\rho = -0.5$ | 7.61***/<4** | 10.35**/5.65*** | 6.96***/<4** | 12.50***/<4*** |
| $\rho = 0$ | 7.53***/<4** | 12.39**/6.51*** | 7.42***/<4*** | 14.40***/<4*** |

Each cell reports PIESS values for predictive $R^2$ for the phylogenetically dispersed subset and the phylogenetically clustered subset, respectively, in the format dispersed/clustered. Asterisks indicate empirical one-sided significance relative to random subsets generated from the same candidate pool and with the same target subset size: \* $p < 0.05$ , \*\* $p < 0.01$ , and \*\*\* $p < 0.001$ . For dispersed subsets, significance was evaluated in the direction of higher PIESS than random subsets; for clustered subsets, significance was evaluated in the direction of lower PIESS than random subsets. In Panel A, $\lambda = 0$ represents the limiting case in which all off-diagonal phylogenetic covariance is removed; under this condition, PIESS is expected to be close to the nominal subset size, with small deviations possible because PIESS is estimated from finite simulation replicates and interpolation against an empirical benchmark curve. Parameter definitions and interpretations are provided in Supplementary Methods Section S4.

The response of PIESS to covariance strength was broadly consistent with the MIESS-based results. Under *λ*-transformed BM covariance structures, PIESS generally increased as *λ* decreased, reflecting the progressive weakening of off-diagonal phylogenetic covariance. At *λ* = 0, where the phylogenetic covariance structure is removed, PIESS values were generally close to the nominal subset size, and neither dispersed nor clustered subsets showed a consistent difference from random subsets. This result is consistent with the expectation that, in the absence of phylogenetic dependence, the selected dispersed and clustered subsets should no longer differ substantially from random subsets in independence-equivalent predictive information.

However, PIESS also showed minor numerical variation that was not present in MIESS because PIESS was estimated from prediction-metric uncertainty under a finite number of simulation replicates rather than from a fully deterministic matrix-based calculation. This explains small discrepancies among repeated evaluations of the same nominal setting. For example, for the dispersed subset with *N* = 512, *s* = 64, and *λ* = 1, the predictive *R*²PIESS was 14.36 in Figure 3c and in Table 2, but 14.59 in the alternative-covariance sensitivity analysis, a difference of approximately 1.6%. The same simulation-based estimation also explains occasional PIESS values slightly exceeding the nominal subset size under the no-phylogenetic-dependence condition. For instance, at *N* = 128 and *s* = 64, the dispersed subset produced PIESS values of 67.41 for MAE when *λ* = 0. Finally, PIESS did not always change monotonically with covariance strength. At *N* = 512 and *s* = 8, the PIESS for predictive *R*²at *λ* = 0.75 that exceeded those at *λ* = 0.25 (7.59 vs. 7.56). These local deviations likely reflect finite simulation variation and metric-specific calibration effects, reinforcing that PIESS should be interpreted primarily by its broad qualitative trends rather than exact parameter-by-parameter monotonicity.

Together, the idealized-tree examples and nested Cricetidae analyses show that the proposed framework can distinguish low-dependence dispersed subsets from random and high-dependence clustered subsets across contrasting tree geometries, candidate-pool sizes, subset sizes, and covariance assumptions. At the same time, they show that the absolute amount of independence-equivalent information in a selected subset should not be interpreted as an intrinsic property of the subset alone. Instead, it depends jointly on the phylogenetic positions and spacing of the selected species, the structure of the underlying phylogeny, the candidate pool from which they were drawn, the target subset size, and the covariance model used to translate tree structure into expected dependence.

## Discussion

Species-level predictive studies often describe evaluation datasets by their nominal number of species, but shared evolutionary history means that species do not necessarily contribute independent information in proportion to their count. In this study, phylogenetically dispersed subsetting was used as a deliberately favorable case for reducing within-subset dependence: if species-level validation sets can approach independence through subset choice alone, dispersed subsets should provide one of the most favorable settings available under fixed subset-size and candidate-pool constraints. Our results show that dispersion did reduce dependence relative to random and clustered subsets, but substantial tree-induced dependence often remained. Thus, even apparently large and carefully selected validation subsets can contain much less effective information than their raw species counts suggest. These findings shift the interpretation of species-level validation sets from a simple counting problem to an information-content problem: the central question is not only how many species are included, but how much independent information those species provide under the evolutionary structure of the dataset.

### Limited Information in Validation Subsets

Under a BM model, a dispersed 64-species subset selected from the 512-species Cricetidae candidate pool yielded a MIESS of only 8.91, and its PIESS estimates for RMSE, MAE, and *R*²were all <15. This result provides a striking quantitative caution for machine-learning applications and is consistent with the broader idea that phylogenetically structured samples may contain substantially less independent information than their nominal sample size suggests (Ané2008, Faes et al. 2009, Bartoszek 2016, Gardner and Organ 2021). The situation becomes even more severe for phylogenetically clustered subsets, which can resemble evaluation sets that inadvertently overrepresent particular clades. For the clustered reference subset selected from the same 512-species Cricetidae candidate pool, MIESS dropped to 1.29, and PIESS estimates were <4. Thus, under the same nominal 64-species evaluation size, clustered sampling provided only a very small fraction of the independent information implied by the species count. Although MIESS and PIESS quantify different targets, their results therefore converged on the same qualitative conclusion: species counts alone can substantially overstate the amount of independent information available for model evaluation.

This limitation has direct implications for model validation and hyperparameter optimization. Standard machine-learning workflows use validation sets or cross-validation to compare parameter configurations and select models expected to generalize beyond the training data. However, when data contain spatial, temporal, hierarchical, or phylogenetic structure, random or unstructured validation strategies can give misleading estimates of model performance, motivating blocked or structure-aware cross-validation strategies (Roberts et al. 2017). Our results extend this concern to the internal information content of the evaluation subset itself: although a nominal validation set of 64 species might appear moderately sized under ordinary independent-sample intuition, it should not automatically be interpreted as a large independent validation sample. In the BM-based Cricetidae analysis, even a dispersed 64-species subset retained far less independence-equivalent information than its nominal species count implied, and clustered subsets could approach the information content of only a few independent observations. Such low effective information can make model selection highly sensitive to noise and to the particular phylogenetic topology represented in the validation set.

This concern follows from the finite-sample problem emphasized by Cawley and Talbot (2010): when a model-selection criterion is estimated from a limited sample, its variance can cause overfitting at the model-selection stage itself and bias subsequent performance evaluation. Such phylogenetically induced information loss can inflate the variance of the validation estimate and make hyperparameter choices more likely to track sampling noise or lineage-specific structure than genuinely generalizable biological relationships.

### The Limits of Dispersion and the Role of Evolutionary Context

Phylogenetically dispersed subsetting reduced within-subset dependence, but it did not make selected subsets independent under a strict BM process (Freckleton et al. 2002). This strong-dependence scenario is far from a theoretical curiosity, because strong phylogenetic signals have been reported for many routinely analyzed biological traits. For example, plant seed mass and plant height showed Pagel’s *λ* values of 1.00 and 0.98, respectively, in a subtropical regional flora (Wang et al. 2021). In rodents, basal metabolic rate and body mass also showed strong phylogenetic signals, with *λ* values of 0.99 and 0.98, respectively (Cutrera and Luna 2025). Comparable patterns have been reported in prokaryotes, where growth-temperature traits show high phylogenetic signals, with Pagel’s *λ* values generally above 0.93, while multiple GC-content measures show even stronger signals, often close to or equal to 1.0 (Hu et al. 2022). Bacterial CRISPR-Cas-related traits also show strong phylogenetic signals, with *λ* values ranging from 0.846 to 0.922 (Lan et al. 2022). Under the *λ*-transformed BM analysis, the dispersed 64-species subset yielded PIESS values well below the nominal size under moderate to strong phylogenetic signal. For example, PIESS for *R*² was 21.59 at λ = 0.75. As phylogenetic signal decreased, PIESS increased; at *λ* = 0.50, it reached 31.96, approximately half of the nominal size. These results indicate that even moderate phylogenetic dependence can substantially reduce the effective information content of evaluation subsets, suggesting that effective sample sizes in species-level evaluation may be much smaller than nominal species counts under typical levels of phylogenetic signal.

### Towards Dependence-Aware Model Evaluation

An evaluation set may contain far fewer independent observations than its species count suggests. This issue has long been recognized in comparative statistical analysis, where phylogenetic non-independence reduces effective degrees of freedom and can substantially alter inference in species-level regression settings (Revell 2010, O’Meara 2012, Slater and Pennell 2014). A practical trade-off in species-level machine-learning evaluation is thus that model assessment must balance maximizing sample size with controlling phylogenetic dependence. Using the full candidate pool generally provides more absolute effective information than any subset, including a maximally dispersed one, even when many species are phylogenetically redundant. However, larger evaluation sets may also contain stronger internal dependence, particularly when additional species come from already well-represented clades. By contrast, phylogenetically dispersed subsets reduce internal dependence and increase independence efficiency, but cannot exceed the total information available in the full pool. Maximizing total effective information and minimizing within-subset dependence are therefore related but distinct goals.

This distinction clarifies the role of phylogenetically dispersed subsetting. Instead of interpreting dispersed subsets as a recommendation to discard available species, they are most useful when the candidate pool is sufficiently large to permit selective sampling and a fixed-size benchmark subset is required, when balanced comparisons across models or datasets are needed, or when a deliberately low-dependence evaluation set is desired. In these contexts, dispersed subsetting provides a reference for assessing how much internal dependence remains after phylogenetic redundancy has been reduced, given a fixed subset size and candidate pool.

Importantly, the trade-off between maximizing total effective information and minimizing within-subset dependence should be interpreted in conjunction with the sensitivity of performance metrics to phylogenetic structure. Beyond reductions in effective sample size, phylogenetic dependence may also induce systematic distortions in performance estimates, leading to potentially over-optimistic evaluations when such structure is not accounted for (Roberts et al. 2017). In cases where phylogenetically adjusted or bias-corrected performance measures are available, or where the metric itself is relatively insensitive to phylogenetic structure, maximizing total effective information using the full candidate pool may be preferable. By contrast, when performance metrics are known to be sensitive to phylogenetic structure and no satisfactory correction is available, reducing within-subset dependence through phylogenetically dispersed sampling becomes a necessary strategy to mitigate evaluation bias.

A comprehensive assessment of metric behavior under phylogenetic dependence would require systematic evaluation across a wider range of predictive performance measures, including commonly used forecasting and regression-based accuracy metrics (Hyndman and Koehler 2006, Jadon et al. 2024). Developing alternative evaluation frameworks that explicitly account for structured dependence remains an important direction for future methodological work.

For regression-type performance summaries such as *R*², one practical extension is to compare observed and predicted values using PGLS (Grafen 1989, Pagel 1999, Symonds and Blomberg 2014), or to use *R*²definitions for correlated data as developed by Ives (2018). Rather than relying solely on an ordinary regression between observed and predicted values, one can compare ordinary *R*²with a phylogenetically adjusted version that incorporates the covariance structure of the evaluation species. Such comparisons do not decompose predictive performance into “feature learning” and “phylogenetic interpolation,” but a substantial reduction after phylogenetic adjustment would indicate that apparent model performance is partly structured by shared evolutionary history within the evaluation set.

Overall, dependence-aware model evaluation should distinguish among three related but distinct objectives: maximizing effective information, accounting for internal dependence within evaluation sets, and interpreting predictive performance in the presence of phylogenetic structure. Phylogenetic dependence cannot always be eliminated through subset design alone, but it can be reduced where possible, quantified explicitly, and incorporated into sensitivity analyses. In this sense, dependence-aware evaluation provides a practical compromise between independence assumptions and data-structural realism, by making the implications of phylogenetic covariance explicit in species-level model assessment. Similar concerns have long been raised in spatial and spatiotemporal modeling, where unaccounted spatial autocorrelation can lead to substantial overoptimism under standard cross-validation procedures (Ploton et al. 2020).

## Funding

This work was supported by remaining funds from the National Natural Science Foundation of China (grant number 31671321). No dedicated funding was received for this specific project.

## Acknowledgments

Many sentences in this article were drafted or refined with the assistance of ChatGPT and Gemini, large language models developed by OpenAI and Google, respectively. The authors thoroughly reviewed and revised each sentence and take full responsibility for the language and content of the entire article.

## Supplementary Materials

Supplementary materials, including Supplementary Methods and Results including Figure S1-S2 and Tables S1–S12, are submitted together with the main text.

## Data availability

Species lists and subset memberships used in the analyses are provided in the Supplementary Materials. The proposed framework was implemented in the R package PhyloTestESS, freely available at https://github.com/HuangR-etc/PhyloTestESS, which includes functions for subset selection, dependence-diagnostic evaluation, and calculation of both MIESS and PIESS to quantify internal phylogenetic dependence and uncertainty in predictive performance metrics.

## Notes

### Competing Interest Statement

The authors have declared no competing interest.

### Summary of Updates

In this revised version, we have introduced a new analytical component, PIESS (prediction-metric-based independence-equivalent sample size), which extends the original framework by quantifying effective information in predictive evaluation metrics (RMSE, MAE, and R2) under phylogenetic dependence. PIESS provides a direct mapping between uncertainty in predictive performance and an independence-equivalent sample size scale, complementing the previously defined MIESS framework. This addition enables a unified assessment of how phylogenetic structure affects both statistical estimation and predictive evaluation. In addition, the manuscript has been structurally revised to improve clarity and readability. We have reorganized several sections of the main text, expanded explanatory content, and added multiple figures and tables to better illustrate key concepts and results. The Supplementary Materials have also been substantially expanded to provide additional methodological detail and supporting results, facilitating interpretation and reproducibility for readers.

