## Supplementary material for "Phylogenetic Dependence and Effective Information in Species-Level Model Evaluation": Supplemementary materials

### Supplementary Materials

Table S5. MeanOffCor in the nested Cricetidae sensitivity analysis. 26
Table S6. MaxOffCor in the nested Cricetidae sensitivity analysis. 26
Table S7. PIESS for RMSE in the nested Cricetidae sensitivity analysis. 27
Table S8. PIESS for MAE in the nested Cricetidae sensitivity analysis. 27
Table S9. MeanOffCor under alternative covariance assumptions in the nested Cricetidae sensitivity analysis. 28
Table S10. MaxOffCor under alternative covariance assumptions in the nested Cricetidae sensitivity analysis. 29
Table S11. PIESS for RMSE under alternative covariance assumptions in the nested Cricetidae sensitivity analysis. 30
Table S12. PIESS for MAE under alternative covariance assumptions in the nested Cricetidae sensitivity analysis. 31
References for Supplementary Materials 32

### Supplementary Methods

#### Section S1. Distance-Based Construction of Dispersed and Clustered Subsets

##### Distance-based descriptors of within-subset phylogenetic structure

Let *C* denote the candidate species pool and *s* the target subset size. For each candidate pool, patristic distances among all species were represented by a distance matrix

$$D = d_{ij},$$

where $d_{ij}$ is the total branch length connecting species *i* and species *j*. For a target subset $S \subset C$ of size *s*, we used distance-based descriptors to quantify different aspects of within-subset phylogenetic dispersion. The first descriptor was the minimum pairwise patristic distance,

$$\text{MinPD}(S) = \min_{i,j \in S; i<j} d_{ij},$$

which reflects the smallest pairwise distance among selected species and therefore acts as a lower-bound constraint on within-subset phylogenetic spacing, preventing the inclusion of near-duplicate or overly clustered species. The second descriptor was the mean pairwise patristic distance,

$$\text{MeanPD}(S) = \frac{2}{s(s-1)} \sum_{i,j \in S; i<j} d_{ij},$$

which summarizes the overall phylogenetic spread of the selected subset. The third descriptor was the mean nearest-neighbor phylogenetic distance,

$$\text{MeanNND}(S) = \frac{1}{s} \sum_{i \in S} \min_{j \in S; j \neq i} d_{ij},$$

which measures the average distance from each selected species to its nearest selected neighbor and therefore reflects local spacing around selected tips. For clustered subset construction, we also used the maximum pairwise patristic distance,

$$\text{MaxPD}(S) = \max_{i,j \in S,, i<j} d_{ij},$$

which represents the greatest phylogenetic separation between any two selected species in the subset. In the clustered-subset context, MaxPD served as a boundary-distance criterion, helping to ensure that even the most distant pair of selected species remained phylogenetically close. Together, these descriptors provide complementary summaries of within-subset phylogenetic structure, capturing local spacing, overall dispersion, and boundary extent at different scales.

##### Construction of phylogenetically dispersed subsets

Phylogenetically dispersed subset construction was formulated as a multi-criterion optimization problem. The primary objective was to maximize MinPD, with MeanPD and MeanNND used as secondary and tertiary criteria, respectively. Thus, between two candidate subsets of the same size, the preferred subset was the one with the larger MinPD; if MinPD was tied, the subset with the larger MeanPD was preferred; if both MinPD and MeanPD were tied, the subset with the larger MeanNND was preferred. This ordered criterion was used to prioritize avoidance of very close species pairs while still favoring broad overall spread and local separation across the candidate-pool tree.

Because exhaustive search over all possible subsets was computationally impractical for large candidate pools, dispersed subsets were constructed using a two-stage heuristic procedure. In the first stage, an initial subset was built by greedy forward selection. The first species was chosen as the species with the largest average patristic distance to all other species in the candidate pool, thereby placing the initial subset at a phylogenetically peripheral position rather than at an arbitrary starting point. Species were then added sequentially until the target subset size was reached. At each step, every unselected candidate was temporarily added to the current subset, and the resulting candidate subset was evaluated using the ordered criteria: larger MinPD, then larger MeanPD, then larger MeanNND. The candidate producing the best ordered value was retained.

In the second stage, the complete greedy subset was refined by one-for-one local swaps. For every species already included in this subset, we considered replacing it with each species not included in the subset, and each resulting one-for-one replacement was re-evaluated under the same ordered criteria. A replacement was accepted if it improved the ordered objective. This procedure was repeated until no further improving exchange could be found.

The greedy initialization was used only to provide a deterministic starting point; the subsequent swap-based refinement allowed the final subset to move away from the initial greedy path when alternative combinations better satisfied the ordered dispersion criteria. Although this heuristic does not guarantee a global optimum, it provides a transparent and computationally practical procedure for constructing strongly dispersed subsets from large candidate pools.

##### Construction of phylogenetically clustered subsets

Phylogenetically clustered subsets were constructed as high-dependence contrast sets rather than as a primary optimization target of the framework. Because clustered subset construction is inherently tree-local rather than globally dispersive, a separate clustering-oriented heuristic was used instead of reversing the dispersed-subset procedure.

The main clustered-subset algorithm is based on a multi-start, global ranking, and exchange-refinement procedure. For each candidate pool and target subset size, each species was in turn used as a starting point, and an initial subset was constructed sequentially by adding species that minimized an ordered clustering criterion. At each step, MeanPD was used as the primary criterion, followed by MeanNND and MaxPD as secondary and tertiary criteria, respectively. This process produced multiple candidate subsets originating from different starting points. Duplicate subsets were removed prior to ranking, and the remaining subsets were compared using the same ordered criterion. The highest-ranked subset was retained as the final clustered subset for each candidate pool and subset size. A subsequent one-for-one exchange procedure between selected and unselected species was then applied using the same ordered criterion to refine subset composition until no further improvement was possible.

**An alternative implementation** based on a nearest-neighbor greedy heuristic was also considered. In this approach, each species in the candidate pool was used as a starting point, and subsets were constructed by iteratively adding the $s-1$ species with the smallest patristic distances to the starting species. Unlike the main procedure, this approach does not perform multi-criteria ranking or exchange-based refinement, and therefore represents a purely local construction strategy based on nearest-neighbor expansion.

This greedy nearest-neighbor approach is computationally simpler because it avoids repeated evaluation of full within-subset distance structure. In relatively balanced trees it often produces similar clustered subsets to the full multi-start plus exchange procedure, but differences may arise under uneven phylogenetic structure or when multiple local clusters compete. For this reason, it was retained as a computationally efficient approximation but was not used in the main analyses reported here.

#### Section S2. Correlation-Based Dependence Diagnostics and MIESS

In the main text, we evaluated within-subset phylogenetic dependence using three diagnostics derived from the model-implied correlation matrix of each selected subset: the mean off-diagonal correlation (MeanOffCor), the maximum off-diagonal correlation (MaxOffCor), and the mean-based independence-equivalent sample size (MIESS). This appendix explains the logic behind these quantities, with particular emphasis on why MIESS can be interpreted as a mean-based effective sample size. The goal here is not to give the most compact mathematical derivation, but to make the logic transparent for readers who may not work with generalized least squares, phylogenetic covariance matrices, or effective sample size on a regular basis.

##### S2.1. From a phylogenetic covariance to a phylogenetic correlation

For a selected subset $S$ containing $s$ species, let $V_S$ denote the phylogenetic covariance matrix among the species in $S$ under a specified evolutionary covariance model. The entries of $V_S$ describe the covariance expected between species because of shared evolutionary history.

Under Brownian motion (BM), for example, the covariance between two species is determined by their shared ancestral branch length. Species that share more recent evolutionary history generally have larger covariance, whereas species that diverged earlier generally have smaller covariance. Other evolutionary models, such as $\lambda$-transformed BM, Ornstein–Uhlenbeck (OU), and Early-burst (EB) models, imply different covariance matrices, but the same diagnostic framework can be applied once $V_S$ has been obtained.

Because the absolute values in a covariance matrix depend on marginal variances as well as dependence, we standardized $V_S$ to the corresponding correlation matrix. Let

$$\Delta = \mathrm{diag}(V_S)$$

be the diagonal matrix containing the marginal variances in $V_S$. The corresponding phylogenetic correlation matrix ($R_S$) is

$$R_S = \Delta^{-1/2} V_S \Delta^{-1/2}.$$

Equivalently, the $(i, j)$-th entry of $R_S$ is

$$(R_S)ij = \frac{(V_S)ij}{\sqrt{(V_S)ii(V_S)jj}}.$$

The diagonal entries of $R_S$ are therefore equal to 1, and the off-diagonal entries describe the pairwise correlations among species induced by the assumed phylogenetic covariance model. These off-diagonal entries are the direct basis of the correlation-based dependence diagnostics.

##### S2.2. Mean and maximum off-diagonal correlations

The first correlation-based diagnostic is the mean off-diagonal correlation,

$$\text{MeanOffCor} = \frac{2}{s(s-1)} \sum_{i<j} (R_S)_{ij}.$$

There are $s(s-1)/2$ unique species pairs in a subset of size $s$. MeanOffCor is therefore the arithmetic mean of all unique pairwise correlations among species in the subset. It summarizes the average level of model-implied dependence within the subset. Lower MeanOffCor indicates weaker average tree-induced dependence.

The second diagnostic is the maximum off-diagonal correlation,

$$\text{MaxOffCor} = \max_{\boldsymbol{i<j}} (\boldsymbol{R_S})_{\boldsymbol{ij}}.$$

MaxOffCor captures the strongest remaining pairwise dependence in the subset. This is useful because a subset may have low average dependence while still containing one pair of closely related species. MeanOffCor and MaxOffCor therefore describe complementary aspects of within-subset dependence: the first summarizes average pairwise dependence, whereas the second captures the strongest local redundancy.

For phylogenetically dispersed subsets, lower MeanOffCor and lower MaxOffCor indicate that the selected species are less dependent under the assumed covariance model. For phylogenetically clustered subsets, higher MeanOffCor and higher MaxOffCor indicate stronger within-subset dependence.

##### S2.3. What question is MIESS trying to answer?

At first glance, one might say that a selected subset containing $s$ species has sample size $s$, because there are $s$ observed species. That is true if we are only counting observations. However, it is not necessarily true if we are asking how much independent information those observations contain. The reason is that species within a phylogenetically structured subset are typically not independent. If two species are closely related, their values may contain partly overlapping information. In that case, counting both species as if they contributed two fully independent pieces of information can be misleading.

The idea of an effective sample size is to answer a more refined question: If these correlated observations were replaced by independent observations, how many independent observations would provide the same amount of information for the inferential target we care about?

In this section, the inferential target is an aggregated mean, leading to the mean-based independence-equivalent sample size (MIESS). This quantity measures the amount of independent information available for estimating an aggregated mean under the assumed correlation structure. It is therefore distinct from the prediction-metric-based independence-equivalent sample size (PIESS), and should not be interpreted as a universal effective sample size for every parameter or statistical task.

For a selected subset, MIESS is calculated as

$$\mathrm{MIESS} = \mathbf{1}^T R_S^{-1} \mathbf{1}.$$

Here, $R_S$ is the correlation matrix of the selected subset, and $\mathbf{1}$ is a column vector of ones of length $s$.

##### S2.4. The reference model for interpreting MIESS

To make the interpretation concrete, suppose that a hypothetical trait is measured on the selected species and that the resulting values are collected in the vector

$$y = (y_1, y_2, \dots, y_s)^T.$$

This trait vector is introduced only to define an inferential target through which the information content of the subset can be interpreted. The calculation of MIESS itself does not require observed trait values at the terminal nodes; it depends only on the assumed within-subset correlation matrix $R_S$.

We consider the simplest possible model for an overall mean:

$$y = \mu 1 + \varepsilon,$$

where

$$1 = (1,1,\dots,1)^T$$

is a column vector of ones of length $s$, $\mu$ is the overall mean, and $\varepsilon$ is the error vector.

We assume that the error vector has covariance structure

$$\mathrm{Var}(\varepsilon) = \sigma^2 R_S.$$

This expression is worth unpacking carefully. First, ε is a vector, not a single random variable. Therefore, Var(ε) is a covariance matrix rather than a single number. Second, $\sigma^2$ is a variance scale parameter that controls the overall magnitude of variation. Third, $R_S$ is the correlation matrix for the selected subset. Because it is a correlation matrix, its diagonal entries are 1, and its off-diagonal entries describe the correlation between different observations in the subset. Thus, $\sigma^2 R_S$ means that all observations share the same marginal variance scale $\sigma^2$, while their dependence structure is described by $R_S$.

##### S2.5. Independent-sample variance scaling as a reference baseline

If observations are independent and each has variance $\sigma^2$, then the variability of the sample mean decreases predictably with sample size. This provides the fundamental reference case for all effective sample size constructions.

In the independent case, adding one additional observation contributes a fixed and non-redundant amount of information, leading to the well-known $1/n$ scaling of the mean variance. This relationship establishes the baseline against which correlated cases must be compared: when observations are not independent, the nominal sample size $n$ no longer directly reflects the amount of independent information available.

This motivates the notion of an effective sample size, which replaces the nominal count of observations with an independence-equivalent quantity that preserves the same inferential precision for the target statistic.

##### S2.6. Variance of the sample mean under independence

Suppose $z_1, z_2, \dots, z_n$ are independent observations, each with variance

$$\mathrm{Var}(z_i) = \sigma^2.$$

Their sample mean is

$$\bar{z} = \frac{1}{n}\sum_{i=1}^{n} z_i.$$

We now compute the variance of $\bar{z}$, the sample mean, step by step:

$$\mathrm{Var}(\bar{z}) = \mathrm{Var}\left(\frac{1}{n}\sum_{i=1}^{n} z_i\right) = \frac{1}{n^2}\mathrm{Var}\left(\sum_{i=1}^{n} z_i\right).$$

Because the $z_i$ are independent, the variance of their sum is the sum of their variances:

$$\mathrm{Var}\left(\sum_{i=1}^{n} z_i\right) = \sum_{i=1}^{n} \mathrm{Var}\,(z_i) = \sum_{i=1}^{n} \sigma^2 = n\sigma^2.$$

Substituting this back gives

$$\mathrm{Var}(\bar{z}) = \frac{1}{n^2} n\sigma^2 = \frac{\sigma^2}{n}.$$

Thus, under independence, the variance of the sample mean decreases at rate $1/n$, providing the baseline scaling law used in the definition of MIESS.

##### S2.7. What estimator should be used when observations are correlated?

Now return to our model

$$\mathbf{y} = \mu\mathbf{1} + \boldsymbol{\varepsilon}, \qquad \mathrm{Var}(\boldsymbol{\varepsilon}) = \sigma^2 R_S.$$

When observations are correlated, the ordinary independent-errors formula is no longer the natural one to use. The standard result in this setting is the generalized least-squares (GLS) result classically associated with Aitken (1936). In modern notation, for the general linear model

$$\mathbf{y} = X\boldsymbol{\beta} + \boldsymbol{\varepsilon}, \qquad \mathrm{Var}(\boldsymbol{\varepsilon}) = \sigma^2 V,$$

the coefficient-estimator form of the Aitken generalized least-squares estimator is

$$\widehat{\boldsymbol{\beta}}_{\mathrm{GLS}} = (X^T V^{-1} X)^{-1} X^T V^{-1} \mathbf{y},$$

Because Aitken's original notation differs from modern linear-model notation, and because his displayed transformation corresponds to the fitted values and therefore includes an additional left multiplication by the design matrix, we use here the standard modern coefficient-estimator form. Its variance-covariance matrix is

$$\mathrm{Var}(\widehat{\boldsymbol{\beta}}_{\mathrm{GLS}}) = \sigma^2 (X^T V^{-1} X)^{-1}.$$

Here, β denotes a vector of model coefficients in the general GLS formula. In the intercept-only model used for MIESS, there is only one coefficient, and this coefficient is the overall mean μ, not a slope. Therefore, for our model,

$$X = \mathbf{1}, \qquad V = R_S.$$

Substituting these into the general GLS formula gives

$$\hat{\mu}_{\mathrm{GLS}} = (\mathbf{1}^T R_S^{-1} \mathbf{1})^{-1} \mathbf{1}^T R_S^{-1} \mathbf{y},$$

with variance

$$\mathrm{Var}(\hat{\mu}_{\mathrm{GLS}}) = \sigma^2 (\mathbf{1}^T R_S^{-1} \mathbf{1})^{-1}.$$

Because this is a scalar, we can rewrite it as

$$\mathrm{Var}(\hat{\mu}_{\mathrm{GLS}}) = \frac{\sigma^2}{\mathbf{1}^T R_S^{-1} \mathbf{1}}.$$

This is the key variance in our derivation.

##### S2.8. What does this GLS variance mean?

The estimator $\hat{\mu}_{\mathrm{GLS}}$ is the generalized least-squares estimator of the overall mean under the assumed correlation structure $R_S$. Its variance tells us how precisely the overall mean can be estimated from this correlated subset. A smaller variance means that the mean is estimated more precisely. A larger variance means that the estimate is less precise. Therefore, in this setting, the quantity

$$\mathbf{1}^T R_S^{-1} \mathbf{1}$$

directly determines the variance of the GLS estimator of the mean and thus acts as an effective sample size. This does not mean that $\mathbf{1}^T R_S^{-1} \mathbf{1}$ is a universal measure of information for every possible inferential task. It is tied specifically to the overall-mean problem considered here. That is why we call the resulting quantity a mean-based independence-equivalent sample size.

##### S2.9. Matching the correlated-sample variance to the independent-sample variance

We now come to the sentence that motivates the definition. From the correlated sample, we have the GLS variance

$$\mathrm{Var}(\hat{\mu}_{\mathrm{GLS}}) = \frac{\sigma^2}{\mathbf{1}^T R_S^{-1} \mathbf{1}}.$$

From an independent sample of size $n$, we know that the variance of the sample mean is

$$\frac{\sigma^2}{n}.$$

Now imagine a hypothetical independent sample of size $n_{\text{eff}}$. If we want that hypothetical independent sample to estimate a mean with exactly the same precision as the actual correlated sample, then the two variances should be equal. That is,

$$\frac{\sigma^2}{\mathbf{1}^T R_S^{-1} \mathbf{1}} = \frac{\sigma^2}{n_{\text{eff}}}.$$

Because the numerators are the same, matching the two variances requires the denominators to be equal:

$$n_{\text{eff}} = \mathbf{1}^T R_S^{-1} \mathbf{1}.$$

Thus, the mean-based independence-equivalent sample size is

$$\text{MIESS} = \mathbf{1}^T R_S^{-1} \mathbf{1}.$$

##### S2.10. Why this can be called an independence-equivalent sample size

The logic is now straightforward. An independent sample of size MIESS would have sample-mean variance

$$\frac{\sigma^2}{MIESS},$$

which is exactly equal to the GLS variance of the mean estimated from the *s* nominal observations in the correlated subset under the correlation structure $R_S$. Therefore, for the purpose of estimating an overall mean, the actual correlated subset the *s* nominal observations behaves as if it were an independent sample of size MIESS.

This is why the quantity can be interpreted as an independence-equivalent sample size. It is important to be precise here. MIESS is not the literal number of observations left after somehow "removing" dependence. No observations are removed. Instead, MIESS tells us how much independent information the correlated sample contains for the specific task of estimating an overall mean. This interpretation follows the general logic of effective sample size (Faes et al. 2009) as the size of an equivalent independent sample for a given inferential objective.

##### S2.11. Why we call it mean-based?

The phrase "mean-based" is important and should not be omitted. Everything above is built around one particular inferential target: the overall mean $\mu$. We used the GLS variance for estimating $\mu$ and matched it to the independent-sample variance of a sample mean. Therefore, the resulting effective sample size is specific to this mean-estimation problem.

This does not mean that our analysis is focused on empirical trait means. In our framework, MIESS is calculated from the assumed within-subset correlation matrix $R_S$, not from observed terminal-node trait values. The mean-estimation problem should instead be understood as a simple reference task: if a hypothetical trait measured on the selected species followed the assumed correlation structure, how many independent observations would provide the same precision for estimating its overall mean?

We use the mean as this reference target because it is simple, interpretable, and relatively model-agnostic. Estimating an overall mean requires no predictor variable, no target regression coefficient, no classification threshold, and no loss function. As a result, MIESS mainly summarizes the information loss caused by within-subset correlation, rather than the details of a particular downstream analysis.

If the inferential target were different, such as a regression slope, a contrast, predictive performance, or a classification metric, the relevant variance or uncertainty expression could be different, and the corresponding effective sample size could also be different. For that reason, it would be misleading to present

$$\mathrm{MIESS} = \mathbf{1}^T R_S^{-1} \mathbf{1}$$

as if it were the one and only effective sample size for the subset in every possible sense. This definition is tied to the precision of estimating an overall mean, MIESS should be interpreted as a mean-based independence-equivalent sample size rather than as a universal effective sample size for all possible inferential targets.

##### S2.12. A simple sanity check: the case of independence

A useful check is the special case in which the observations are independent. Then

$$R_S = I,$$

where $I$ is the identity matrix. Since

$$I^{-1} = I,$$

we obtain

$$MIESS = \mathbf{1}^T R_S^{-1} \mathbf{1} = \mathbf{1}^T I^{-1} \mathbf{1} = \mathbf{1}^T I \mathbf{1} = \mathbf{1}^T \mathbf{1}.$$

Because $\mathbf{1}$ has length $s$, the quantity $\mathbf{1}^T \mathbf{1}$ is simply the sum of $s$ ones:

$$\mathbf{1}^T \mathbf{1} = s.$$

So under independence,

$$\mathrm{MIESS} = s.$$

This is exactly what we would want. If the observations are already independent, then the independence-equivalent sample size should equal the nominal subset size.

##### S2.13. Why positive correlation usually reduces MIESS

Although the exact numerical effect depends on the full structure of $R_S$, the intuition is simple. If observations are positively correlated, they tend to move together. That means part of the information in one observation overlaps with information in another observation. In that situation, adding more observations does not increase independent information as efficiently as it would under independence.

As a result, MIESS is often smaller than the literal number of species in the subset. This is not because observations disappear. It is because correlated observations are partly redundant for estimating the mean.

In the extreme case where all species are nearly perfectly correlated, a subset containing many species may behave much more like **a single independent observation** than like multiple independent observations. Conversely, when species are weakly correlated, MIESS approaches **the nominal subset size *s***.

##### S2.14. Relationship among MeanOffCor, MaxOffCor, and MIESS

MeanOffCor, MaxOffCor, and MIESS describe related but non-identical aspects of within-subset dependence.

- MeanOffCor asks: how correlated are species pairs on average?
- MaxOffCor asks: does the subset contain at least one strongly correlated species pair?
- MIESS asks: how many independent observations would provide the same precision for estimating an overall mean under the full correlation structure?

The first two diagnostics are pairwise summaries of the off-diagonal entries of $R_S$. MIESS, by contrast, uses the inverse of the full correlation matrix $R_S^{-1}$. It therefore reflects the joint dependence structure of the subset rather than only the average or maximum pairwise correlation.

These diagnostics are expected to move in consistent directions, but they are not mathematically identical. A subset with low MeanOffCor may still have a high MaxOffCor if it contains one closely related pair. Two subsets with similar MeanOffCor may have different MIESS values if their correlations are distributed differently across the matrix. For this reason, the three diagnostics were used together rather than treating any single number as a complete description of within-subset dependence.

In the main text, lower MeanOffCor, lower MaxOffCor, and higher MIESS are interpreted as evidence of weaker within-subset phylogenetic dependence. For clustered contrast subsets, higher MeanOffCor, higher MaxOffCor, and lower MIESS are interpreted as evidence of stronger within-subset phylogenetic dependence.

##### S2.15. What MIESS is useful for in our setting

In our framework, MIESS is not intended to replace the distance-based subset criteria or the direct correlation-based summaries. Rather, it provides an additional and very interpretable, structure-based summary of the within-subset dependence structure.

Specifically, it answers the following question:

> If a hypothetical trait followed the assumed within-subset correlation structure, and if the goal were to estimate its overall mean from this subset, how many independent observations would provide the same precision as the actual correlated subset?

That is a limited question, but it is also a meaningful and intuitive one. For readers who want a single number with a familiar sample-size interpretation, MIESS can be more accessible than a full matrix summary.

At the same time, MIESS should be interpreted with care. A subset could have the same MIESS as another subset while differing in other aspects of its dependence structure. Therefore, this quantity should be viewed as one interpretable summary, not as a complete replacement for all other summaries.

##### S2.16. Why trait-level phylogenetic signal metrics were not used as subset-dependence diagnostics

Trait-level phylogenetic signal metrics, including Pagel's $\lambda$ and Blomberg's $K$ (Pagel 1999, Blomberg et al. 2003), were not used as criteria for comparing within-subset dependence because they address a different question. These metrics evaluate how strongly a particular observed trait conforms to phylogenetic expectations under a specified model, whereas the diagnostics used here quantify the dependence structure among the species in a selected subset.

This distinction can be illustrated using Pagel's $\lambda$. Suppose a trait is generated exactly under a BM model, corresponding to the $\lambda$ =1 covariance structure on the full tree. If we then examine two different subsets of the same tree, one phylogenetically dispersed and the other phylogenetically clustered, restricting the data to either subset does not change the generating process itself. The expected covariance structure within each subset is still the BM covariance matrix for that subset, corresponding to $\lambda$ = 1. However, their absolute within-subset dependence can be very different. The clustered subset may contain species with high pairwise correlations because they share recent ancestry, whereas the dispersed subset may contain species with much lower pairwise correlations because they are drawn from more distant parts of the tree.

Thus, two subsets can have the same theoretical value of $\lambda$ under the generating model while differing greatly in MeanOffCor, MaxOffCor, and MIESS. Pagel's $\lambda$ describes how the off-diagonal covariance structure of an observed trait compares with a BM expectation on a given tree; it does not directly measure the absolute magnitude of correlation among the species in a subset. Similarly, Blomberg's $K$ evaluates observed trait variation relative to BM expectations, rather than directly summarizing the subset's internal covariance or correlation structure.

This distinction is especially important in the present study because our goal is not to evaluate the phylogenetic signal of a particular trait. Instead, we ask how much internal phylogenetic dependence remains within dispersed, random, and clustered species subsets, and how much effective information those subsets contain under a specified model-implied correlation structure. A subset can therefore have strong tree-induced dependence even when no particular measured trait is being analyzed, and different subsets can differ substantially in their internal dependence even when the same trait-generating model applies to all of them.

For this reason, trait-level phylogenetic signal metrics are not appropriate as direct criteria for comparing the internal dependence of dispersed, random, and clustered subsets. The present study instead uses distance-based criteria for subset construction and model-implied correlation diagnostics, including MeanOffCor, MaxOffCor, and MIESS, to quantify the resulting within-subset dependence.

##### S2.17. A concise summary of the derivation

For readers who want the logic in one place, the derivation of MIESS can be summarized as follows:

1. Assume the intercept-only model

$$\mathbf{y} = \mu\mathbf{1} + \boldsymbol{\varepsilon}, \qquad \mathrm{Var}(\boldsymbol{\varepsilon}) = \sigma^2 R_S.$$

2. Under GLS, the estimator of $\mu$ has variance

$$\mathrm{Var}(\hat{\mu}_{\mathrm{GLS}}) = \frac{\sigma^2}{\mathbf{1}^T R_S^{-1}\mathbf{1}}.$$

3. For $n$ independent observations with common variance $\sigma^2$, the variance of the sample mean is

$$\frac{\sigma^2}{n}.$$

4. Define the mean-based independence-equivalent sample size by requiring these two variances to be equal:

$$\frac{\sigma^2}{\mathbf{1}^T R_S^{-1}\mathbf{1}} = \frac{\sigma^2}{n_{\mathrm{eff}}}.$$

5. Solving gives:

$$\mathrm{MIESS} = \mathbf{1}^T R_S^{-1}\mathbf{1}.$$

MIESS is therefore the size of an independent sample that would provide the same precision for estimating an overall mean as the actual correlated subset under the assumed correlation structure.

#### Section S3. Prediction-Metric-Based Independence-Equivalent Sample Size

MIESS provides a mean-based diagnostic of subset-level information content. Species-level machine-learning evaluation, however, typically focuses on predictive-performance metrics rather than mean estimation. Because effective sample size governs the stability of performance estimates across repeated samples, larger effective sample sizes correspond to narrower 95% empirical intervals. We therefore additionally performed a prediction-metric-based calibration to estimate how many independent test species would yield a comparable level of uncertainty in common predictive metrics.

This analysis was first performed for 64-species subsets selected from the 512-species *Cricetidae* candidate pool. We focused on the phylogenetically dispersed and phylogenetically clustered subsets as contrasting low- and high-dependence cases. The prediction-metric-based independence-equivalent sample size (PIESS) was estimated separately for root mean squared error (RMSE), mean absolute error (MAE), and predictive $R^2$.

For computational efficiency, simulations were conducted directly on the selected subsets rather than on the full 512-species tree. This simplification does not change the simulated distribution for the selected species. Under a multivariate normal model, the marginal distribution of any selected subset of variables is also multivariate normal. The mean vector of that marginal distribution is obtained by retaining the components of the full mean vector for the selected species, and its covariance matrix is obtained by retaining the rows and columns for those same species in the full covariance matrix. Therefore, simulating values on the full 512-tip tree and then extracting the selected 64 species is statistically equivalent, for the present purpose, to simulating them directly using the subset-specific $64 \times 64$ correlation matrix. We therefore first fixed the 64-species dispersed and clustered subsets, extracted their subset-specific BM correlation matrices from the full $512 \times 512$ correlation matrix, and conducted the simulations directly on these $64 \times 64$ matrices.

For each target subset $S$, we denoted its BM correlation matrix as $R_{\mathrm{BM}}(S)$. We then constructed a $\lambda$-transformed correlation matrix as

$$R_{\lambda,ij}(S) = \begin{cases} 1, & i = j, \\ \lambda R_{\mathrm{BM},ij}(S), & i \neq j. \end{cases}$$

Thus, $\lambda = 0$ corresponds to an independent-sample benchmark, whereas $\lambda = 1$ retains the full BM phylogenetic correlation structure within the subset.

For the $\lambda = 1$ phylogenetically structured scenario, we repeated the simulation 10,000 times separately for the dispersed and clustered subsets. Minor non-monotonic patterns across subset sizes may occur due to stochastic variation, reflecting finite-sample variability across simulation replicates under a fixed data-generating process. In each replicate, we generated a true value vector and a prediction-error vector as

$$y \sim \mathcal{N}\big(0, R_{\lambda=1}(S)\big),$$

$$u \sim \mathcal{N}\big(0, \sigma_u^2 R_{\lambda=1}(S)\big),$$

and defined the simulated predicted values as

$$\hat{y} = y + u.$$

Here, $y$ represents the true species-level values, $u$ represents prediction errors, and $\hat{y}$ represents predicted values. The true values and prediction errors were generated independently, but both followed the same subset-specific phylogenetic correlation structure. This design allows prediction errors themselves to contain lineage-specific structure, rather than assuming that errors are independent across species. We set $\sigma_u^2 = 0.1$ so that prediction performance was neither near-perfect nor dominated by error variance. The same error-variance setting was used for all subset types, $\lambda$ conditions, and sensitivity analyses, ensuring that differences among results reflected differences in sample structure and covariance assumptions rather than arbitrary changes in prediction difficulty.

For each replicate and each subset $S$, we calculated RMSE, MAE, and predictive $R^2$ as

$$\mathrm{RMSE}(S) = \sqrt{\frac{1}{s}\sum_{i\in S}(y_i - \hat{y}_i)^2},$$

$$\mathrm{MAE}(S) = \frac{1}{s}\sum_{i\in S}|y_i - \hat{y}_i|,$$

and

$$R^2(S) = 1 - \frac{\sum_{i\in S}(y_i - \hat{y}_i)^2}{\sum_{i\in S}(y_i - \bar{y}_S)^2}.$$

Predictive $R^2$ was not truncated at zero, because negative values indicate that the predictions performed worse than using the subset mean as the prediction.

For each subset and each metric, the 10,000 simulation replicates produced an empirical distribution of the corresponding performance metric. Let $m_1, m_2, \dots, m_B$ denote the simulated values of a given metric $m$, where $m$ is RMSE, MAE, or predictive $R^2$, and $B = 10{,}000$. We summarized the uncertainty of each metric using the width of its 95% empirical interval:

$$W_m(S) = Q_{0.975}(m_1, \dots, m_B) - Q_{0.025}(m_1, \dots, m_B).$$

This interval describes the empirical spread of repeated simulation outcomes and should not be interpreted as a bootstrap confidence interval for a single observed dataset.

To translate the metric uncertainty of a phylogenetically structured 64-species subset into an independence-equivalent sample size, we constructed an independent-sample benchmark under $\lambda = 0$. This benchmark was not generated by repeatedly sampling species from the dispersed or clustered subsets. Once $\lambda = 0$, all off-diagonal correlations are removed, so species identity and subset origin no longer affect the dependence structure. Therefore, for each independent benchmark sample size $n = 4, 5, \dots, 64$, we simulated 10,000 independent samples using an $n \times n$ identity correlation matrix:

$$y \sim \mathcal{N}(0, I_n), \quad u \sim \mathcal{N}(0, \sigma_u^2 I_n),$$

and

$$\hat{y} = y + u.$$

For each $n$, RMSE, MAE, and predictive $R^2$ were calculated across the 10,000 replicates, and their 95% empirical interval widths were obtained. These values formed a common independent-sample benchmark curve for each metric, denoted $W_m^{\mathrm{ind}}(n)$.

For a given 64-species phylogenetic subset $S$ under $\lambda = 1$, we then estimated the metric-specific independence-equivalent sample size by matching its empirical interval width to the corresponding independent benchmark curve:

$$PIESS_m(S) = \arg\min_n \left| W_m^{\mathrm{ind}}(n) - W_m^{\mathrm{phylo}}(S) \right|,$$

where $W_m^{\mathrm{phylo}}(S)$ is the 95% empirical interval width for metric $m$ in subset $S$ under $\lambda = 1$ and $W_m^{\mathrm{ind}}(n)$ is the corresponding interval width for an independent benchmark sample of size $n$.

Because the benchmark curve was evaluated at integer sample sizes, we estimated $PIESS_m(S)$ by linear interpolation along the independent-sample benchmark curve. The dispersed and clustered subsets were matched separately to the same independent benchmark curve, producing metric-specific PIESS estimates for RMSE, MAE, and predictive $R^2$.

When the phylogenetic subset interval width fell between two adjacent independent benchmark values, the independence-equivalent sample size was estimated by linear interpolation along the independent benchmark curve. If the interval width of a phylogenetic subset exceeded the benchmark interval width at the smallest benchmark size, the result was reported as below the lower calibration limit rather than extrapolated. Conversely, if the interval width was narrower than the benchmark interval width at the largest benchmark size, the result was reported as above the upper calibration limit unless an extended benchmark curve was explicitly used. In the main BM PIESS analyses, the independent benchmark was evaluated over $n$ = 4–32. In the alternative-covariance PIESS sensitivity analysis, the benchmark was extended to $n$ = 4–70.

We first conducted sensitivity analyses across multiple combinations of candidate-pool size ($N$) and subset size ($s$) to assess the stability of the main results with respect to sampling design. We additionally performed sensitivity analyses under alternative phylogenetic covariance assumptions. The main prediction-metric-based calibration used the $\lambda = 0$ condition as an independent-sample benchmark and the $\lambda = 1$ BM condition as the primary phylogenetically structured scenario. To examine whether the resulting conclusions depended on this covariance assumption, we repeated the phylogenetically structured simulations for selected representative subsets under $\lambda$-transformed BM, OU, and tree-height-standardized EB covariance structures. For the EB model, branch-specific rates were evaluated on normalized tree time, $\mu = t/H$, before constructing the transformed covariance matrix. RMSE, MAE, and predictive $R^2$ were summarized using the same 95% empirical interval-width procedure. These analyses were interpreted as robustness checks for the qualitative pattern of prediction-metric uncertainty, rather than as a separate definition of effective sample size.

PIESS is specific to the performance metric, simulation setting, covariance structure, and subset being evaluated. It was therefore used as an empirical complement to MIESS, rather than as a universal measure of effective sample size.

The PIESS calculation used 10,000 simulation replicates to estimate the 95% empirical interval width of RMSE, MAE, and predictive $R^2$ for each subset or independent benchmark sample size. This simulation step should be distinguished from the random-subset baseline: 1,000 random subsets were drawn from the same candidate pool, and each was assigned a PIESS value using the same interval-width matching procedure. The resulting 1,000 PIESS values formed the empirical baseline for evaluating whether the selected dispersed or clustered subset was unusually high or low in metric-specific effective information.

#### Section S4. Covariance model parameterization and alternative evolutionary processes

To ensure interpretability and comparability across phylogenetic covariance models, we implemented a unified framework in which all evolutionary models were expressed as modifications of a common BM baseline. In all cases, the starting point is a phylogenetic covariance matrix constructed under a standard BM process on a fixed phylogeny, where covariance between species increases with shared evolutionary history (i.e., shared branch length from the root).

##### S4.1 BM baseline and independence benchmark

Under standard BM, the covariance between any pair of species is proportional to their shared evolutionary history, and the resulting covariance matrix contains non-zero off-diagonal elements that encode phylogenetic dependence.

To provide a common reference point across all models, we define a shared independence benchmark corresponding to a diagonal covariance matrix (i.e., no phylogenetic correlation among species). In the $\lambda$-transformed BM framework, this benchmark is obtained at $\lambda = 0$. All other models, OU and EB, are interpreted relative to this same independence reference to ensure comparability of effective-information measures (MIESS and PIESS) across different evolutionary assumptions.

##### S4.2 λ-transformed BM model

The $\lambda$ transformation provides a simple and widely used way to modulate the strength of phylogenetic signal by rescaling the off-diagonal elements of the BM covariance matrix.

Formally, $\lambda$ acts as a multiplicative factor on all covariances between species:

- When $\lambda = 1$, the original BM covariance structure is recovered.
- When $\lambda = 0$, all off-diagonal elements are removed, producing a diagonal covariance matrix corresponding to phylogenetically independent observations.
- Intermediate values of $\lambda$ ($0 < \lambda < 1$) gradually reduce phylogenetic covariance while preserving marginal variances.

Importantly, $\lambda$ does not change the variance of individual species but only adjusts the strength of shared evolutionary dependence.

##### S4.3 OU model and phylogenetic half-life scaling

The OU model describes evolution under stabilizing selection toward an optimal trait value. In this model, phylogenetic dependence decays over evolutionary time because trait values are pulled toward the optimum.

Instead of using the standard selection strength parameter ($\alpha$) directly, we reparameterize the OU process using the phylogenetic half-life ratio $h/H$, defined as:

- $h$: phylogenetic half-life, defined as the time required for phylogenetic correlation to decay by 50%, given by:

  $$h = \frac{ln(2)}{\alpha},$$

  where $\alpha$ is the strength of stabilizing selection controlling the exponential decay rate of trait covariance.
- $H$: total height of the phylogenetic tree (from root to tips)

The ratio $h/H$ provides a dimensionless measure of how quickly phylogenetic signal decays relative to the depth of the tree:

- Small $h/H$ values correspond to rapid decay of phylogenetic signal. Under this condition, lineages are quickly pulled toward the shared optimum, and tip values can be interpreted as stochastic deviations around this optimum with little covariance attributable to shared ancestry. The resulting correlation structure therefore approaches the independent-sample benchmark.
- Large $h/H$ values correspond to slow decay of phylogenetic signal. In this case, ancestral effects persist for a larger fraction of the tree depth, producing stronger phylogenetic covariance and behavior closer to BM.

This scaling allows OU processes defined on different tree sizes to be directly compared.

##### S4.4 Time-dependent rate transformation and covariance construction

The EB model extends Brownian motion by allowing evolutionary rates to vary through time along the phylogeny. This variation is governed by a dimensionless parameter $\rho$ and implemented on normalized time $\mu = t/H$, where $t$ denotes absolute evolutionary time and $H$ is total tree height. This normalization ensures that $\rho$ is comparable across phylogenies with different absolute depths. EB therefore provides a framework in which branch-length accumulation is later modified through time-dependent rate heterogeneity.

###### Layer 1: Normalized evolutionary time

To ensure comparability across phylogenetic trees with different total depths, we first define a normalized evolutionary time scale. This step is necessary because all subsequent rate modulation and covariance construction depend on a consistent time coordinate system that is invariant to absolute tree height.

We define normalized evolutionary time:

$$\mu = t/H,$$

where:

- $t$: absolute evolutionary time measured from the root,
- $H$: total root-to-tip height of the tree,
- $\mu \in [0, 1]$, with $\mu = 0$ at the root and $\mu = 1$ at the tips.

This normalization ensures that the same value of the model parameter $\rho$ has a consistent interpretation across phylogenetic trees with different absolute timescales. Without this rescaling, the EB rate function defined below would not be directly comparable across datasets with different evolutionary depths.

###### Layer 2: Time-dependent evolutionary rate function

Given the normalized time scale, we define a time-dependent evolutionary rate function that governs how variance accumulates along each branch.

The rate function is specified as:

$$r(\mu) = \exp(\rho\mu)$$

where $\rho \in \mathbb{R}$ controls the direction and magnitude of temporal heterogeneity in rate accumulation.

This formulation allows flexible deviation from the Brownian-motion assumption:

- When $\rho = 0$, the rate is constant over time, recovering the standard BM case.
- When $\rho > 0$, evolutionary rates increase toward the tips, implying accelerated late-stage variance accumulation.
- When $\rho < 0$, evolutionary rates are concentrated deeper in the tree, producing an early-burst pattern.

##### Layer 3: Branch-level variance accumulationType equation here.

We begin by recalling that under Brownian motion, phylogenetic covariance is directly proportional to shared evolutionary history, such that each covariance entry is determined by the depth of the most recent common ancestor (MRCA) of two species. In this formulation, the MRCA depth can be interpreted equivalently as a sum of branch lengths along the shared ancestral path, and thus BM covariance can be viewed as being constructed directly from additive branch-length contributions.

In the EB framework, we retain this same construction principle, but modify the underlying branch lengths through a time-dependent rate function. Specifically, the rate function transforms the original Brownian-motion branch lengths into modified branch lengths, which preserve the same tree topology but alter the amount of variance accumulated along each segment.

Importantly, these modified branch lengths serve as the direct inputs for covariance construction. That is, just as BM covariance entries are given by shared MRCA branch length accumulation, EB covariance entries are obtained by replacing each original branch-length contribution with its transformed counterpart. This defines a one-to-one correspondence between modified branch lengths and the elements of the EB covariance matrix..

For a branch connecting a parent node at time $t_p$ and a child node at time $t_c$, variance accumulation under EB model is defined as:

$$L_\rho(t_p, t_c) = \int_{t_p}^{t_c} r(t/H)dt,$$

which yields:

$$L_\rho(t_p, t_c) = \begin{cases} \frac{H}{\rho}\left[\exp\left(\frac{\rho t_c}{H}\right) - \exp\left(\frac{\rho t_p}{H}\right)\right], & \rho \neq 0 \\ t_c - t_p, & \rho = 0 \end{cases}$$

Thus, Brownian motion emerges as a special case of the EB framework when $\rho$=0, confirming that EB is a generalization rather than a separate model.

This step is the only stage in which the temporal rate function modifies the accumulation of variance; the underlying phylogenetic topology remains unchanged.

##### Layer 4: Covariance construction on the phylogeny

Having defined branch-wise variance accumulation, we construct the phylogenetic covariance structure.

**Brownian-motion baseline**

Under standard BM, covariance between species *i* and *j* is proportional to their shared evolutionary history:

$$V_{ij}^{BM} = t_{MRCA(i,j)}$$

where $t_{MRCA(i,j)}$ is the total branch length from the root to the most recent common ancestor (MRCA) of species *i* and *j*.

This defines the fundamental phylogenetic dependence structure. All subsequent models are constructed as modifications of this shared-path geometry.

**EB-transformed covariance (pre-normalization)**

Under the EB model, the same MRCA-based tree topology is retained, but branch-wise variance accumulation is replaced by the transformed branch lengths derived in Layer 3.

We define:

$$\tilde{V}_{ij}^{EB} = L_\rho(t_{MRCA(i,j)}).$$

This matrix preserves the original phylogenetic topology while altering the temporal distribution of variance accumulation along shared evolutionary paths.

Thus:

- BM assumes linear accumulation of variance over time,
- EB redistributes this accumulation through a nonlinear rate function.

At this stage, $\tilde{V}_{ij}^{EB}$ remains a covariance matrix with marginal variance heterogeneity and is not yet suitable for direct comparison across species or trees.

##### Layer 5: Normalization to correlation space (critical step)

Because EB induces non-stationary variance accumulation across species, all covariance matrices are standardized before downstream analysis:

$$V^{EB} = D^{-1/2}\tilde{V}^{EB}D^{-1/2}$$

where

$$D = diag\left(\tilde{V}^{EB}\right)$$

This transformation ensures that all diagonal elements of $\tilde{V}^{EB}$ equal 1, producing a proper correlation matrix.

This step is essential because:

- EB modifies marginal variances in a time-dependent manner,
- raw covariance values are not directly comparable across different trees or parameter settings,
- downstream dependence diagnostics must reflect correlation structure rather than scale-dependent variance differences.

After this transformation:

- $V^{EB}$ represents pure dependence structure,
- all metrics (MeanOffCor, MaxOffCor, MIESS, PIESS) are computed on standardized space,
- differences in results reflect dependence structure rather than variance scaling artifacts.

**Unified interpretation of notation (explicit clarification)**

To avoid ambiguity, we summarize the role of each matrix:

- $V^{BM}$: original Brownian-motion covariance matrix (baseline model)
- $\tilde{V}^{EB}$: EB-transformed covariance matrix prior to normalization (raw transformed variance structure)
- $V^{EB}$: standardized EB correlation matrix used for all downstream dependence diagnostics

This separation ensures that:

- BM defines the reference evolutionary geometry,
- EB modifies only variance accumulation along branches,
- normalization isolates dependence structure from marginal variance effects.

##### Interpretation of EB parameter *ρ*

| *P* value | r(t)=exp(ρt) behavior (at t=1 intuition) | Interpretation |
|---|---|---|
| 0 | r = 1 | BM baseline (no temporal modulation) |
| small negative (e.g. −0.5) | r < 1 | weak early-burst |
| moderate negative (−1 to −2) | r ≪ 1 | strong early-burst dynamics |
| large negative (≤ −4) | r → 0 | extreme early-burst regime (high temporal heterogeneity) |

The EB parameter ρ controls a monotonic transition from Brownian-motion dynamics ($\rho$ = 0) to progressively stronger early-burst regimes characterized by increasing concentration of evolutionary variance in deeper branches and extreme early-burst regime (high temporal heterogeneity and substantially altered phylogenetic dependence structure after normalization)

#### S4.5 Unified interpretation and dual-benchmark framework

Although $\lambda$-BM, OU, and EB models differ in biological interpretation, all three can be viewed as mechanisms that modulate the strength and structure of phylogenetic covariance relative to a common independence benchmark defined by the $\lambda$ = 0 BM model (diagonal covariance matrix).

Importantly, the $\lambda$ = 0 benchmark is used exclusively to define the scale of effective-information metrics, PIESS. Specifically, PIESS values are constructed by mapping prediction-metric uncertainty to an independence-equivalent sample size using this theoretical reference, and this mapping is applied consistently to all observed subsets (dispersed and clustered) as well as to random subsets.

In contrast, statistical inference for both correlation-based dependence diagnostics and PIESS relied on a second, empirical benchmark based on 1,000 random subsets drawn from the same candidate pool and with the same target subset size. Each random subset was processed in the same way as the selected dispersed and clustered subsets. For the correlation-based diagnostics, this involved calculating MeanOffCor, MaxOffCor, and MIESS. For PIESS, this involved running 10,000 replicate simulations for each random subset, calculating RMSE, MAE, and predictive $R^2$ across replicates, and using the resulting 95% empirical metric width for calibration. These random-subset values provided the null distribution for one-sided significance testing, allowing evaluation of whether the selected dispersed or clustered subsets deviated from expectation under the same phylogenetic and sampling constraints.

Thus, the λ = 0 benchmark defines the absolute scale of independence-equivalent information, whereas the random-subset distribution defines the relative statistical reference. These two benchmarks serve distinct but complementary roles and are not interchangeable.

Across covariance structures, parameter settings were selected to span contrasting dependence regimes rather than a single monotonic continuum. In the $\lambda$-transformed BM model, decreasing $\lambda$ moves the covariance structure toward the independence benchmark. In the OU model, smaller $h/H$ values produce faster decay of phylogenetic dependence and therefore approach the independence benchmark. In the EB model used here, $\rho$ = 0 recovers the BM baseline, whereas increasingly negative $\rho$ values impose stronger root-weighted temporal variance accumulation. These EB settings test robustness to temporal heterogeneity in branch-specific variance accumulation, but they should not be described as near-independence regimes.

### Supplementary Results

#### Section S1. Detailed Distance-Based Validation of Dispersed and Clustered Subsets

To visualize the empirical behavior of the subset-selection procedure, we examined the locations of the selected 64-species dispersed and clustered subsets within the 512-species Cricetidae candidate pool. The dispersed subset was distributed across widely separated regions of the Cricetidae phylogeny, whereas the clustered reference subset was concentrated within a local region of Arvicolinae (Fig. S1). These contrasting spatial patterns motivated the distance-based comparisons with random subsets reported below.

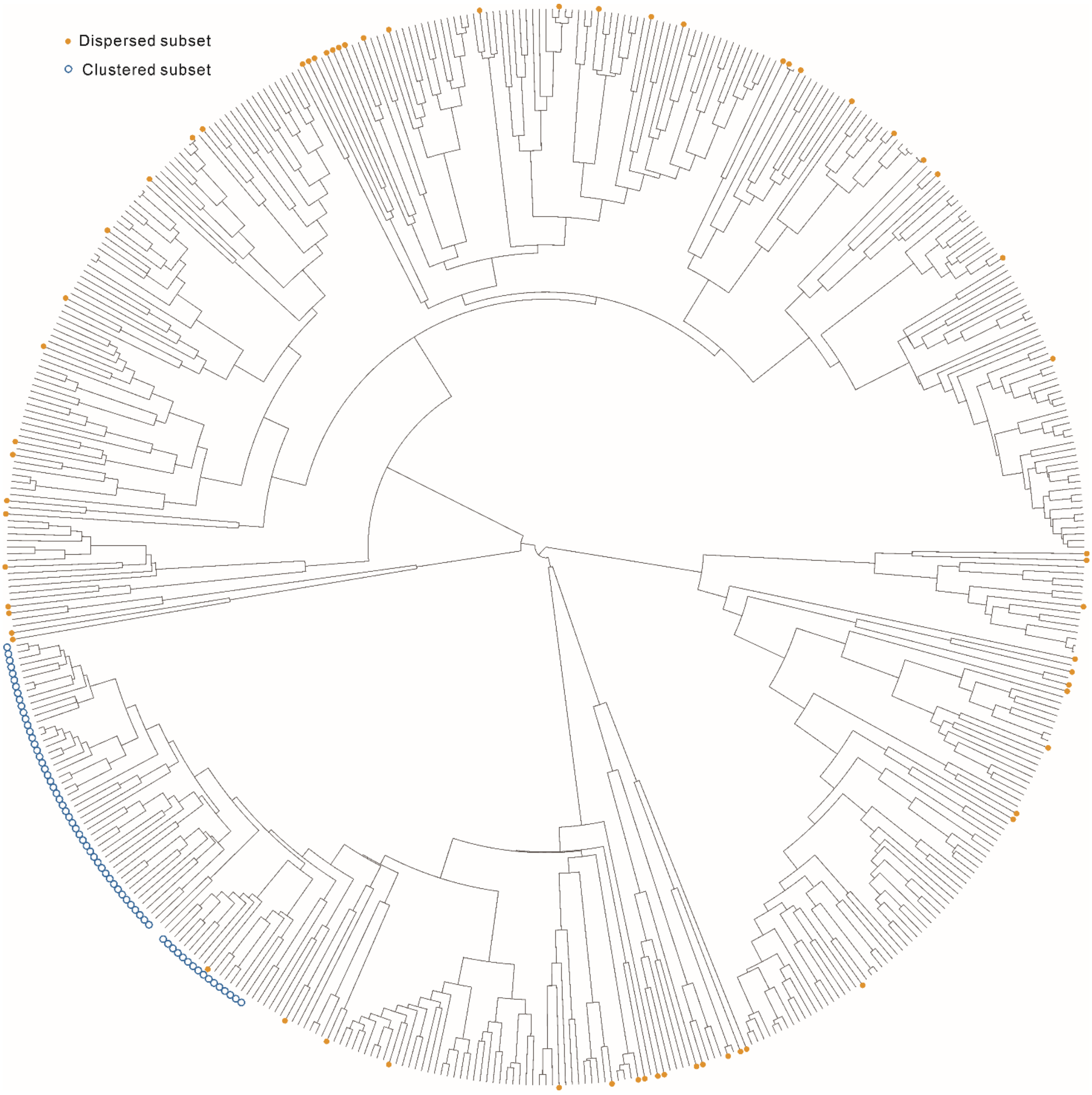

**Figure S1. Phylogenetic distribution of selected Cricetidae subsets.**
The Cricetidae phylogeny was generated by pruning the species-level mammalian phylogeny of Upham et al. (2019). The empirical candidate pool comprised 512 extant cricetid species, from which 64 species were selected for each subset. In the circular phylogeny, orange filled circles indicate species in the phylogenetically dispersed subset, whereas blue open circles indicate species in the phylogenetically clustered reference subset. The dispersed subset is distributed across widely separated regions of the Cricetidae phylogeny, whereas the clustered reference subset is concentrated within a localized region of the same candidate pool, with all selected species belonging to Arvicolinae. Visual proximity among selected tips in the circular layout does not necessarily imply short patristic distance; all distance-based metrics were calculated from branch-length distances on the original tree.

Consistent with this visual contrast, we compared the selected subsets with 1000 random subsets of the same size and asked whether they were displaced toward the expected tails of the random-subset distributions for distance-based criteria (Fig. S2). Dispersion was evaluated using MinPD, MeanPD, and MeanNND, for which larger values indicate stronger phylogenetic dispersion; clustering was evaluated using MaxPD, MeanPD, and MeanNND, for which smaller values indicate stronger phylogenetic clustering. For the phylogenetically dispersed subset, MinPD was 24.18, compared with a random-subset mean of 2.38 and maximum of 8.90, placing the selected dispersed subset beyond all random subsets for this closest-pair criterion (Fig. S2a). The two shared distance metrics showed the same overall contrast. MeanPD was higher in the dispersed subset (64.11; random-subset mean = 61.72; random maximum = 65.04) but much lower in the clustered reference subset (16.85; random minimum = 53.58; Fig. S2b). MeanNND was also higher in the dispersed subset (29.83; random-subset mean = 18.10; random maximum = 22.66) but lower in the clustered reference subset (6.18; random minimum = 13.76; Fig. S2c).

For the boundary criterion of the clustered reference subset, MaxPD was not shown as a separate distribution because all 1000 random subsets had the same MaxPD value of 77.57. In contrast, the clustered reference subset had a much smaller MaxPD of 20.25, indicating that even the most distant pair within this subset remained phylogenetically close relative to random sampling from the same candidate pool.

Relative to the empirical null distributions generated by random sampling, the dispersed subset fell in the upper tail for all three dispersion-oriented criteria, with one-sided empirical $p$-values of 0.001 for MinPD, 0.027 for MeanPD, and 0.001 for MeanNND. The clustered reference subset fell in the lower tail for all three clustering-oriented criteria, with one-sided empirical $p$-values of 0.001 for MaxPD, MeanPD, and MeanNND. Together, these comparisons show that the selected dispersed and clustered subsets occupied opposite ends of the random-subset distributions within the same Cricetidae candidate pool. This confirms that distance-based subset selection behaved as intended in a realistic empirical phylogeny.

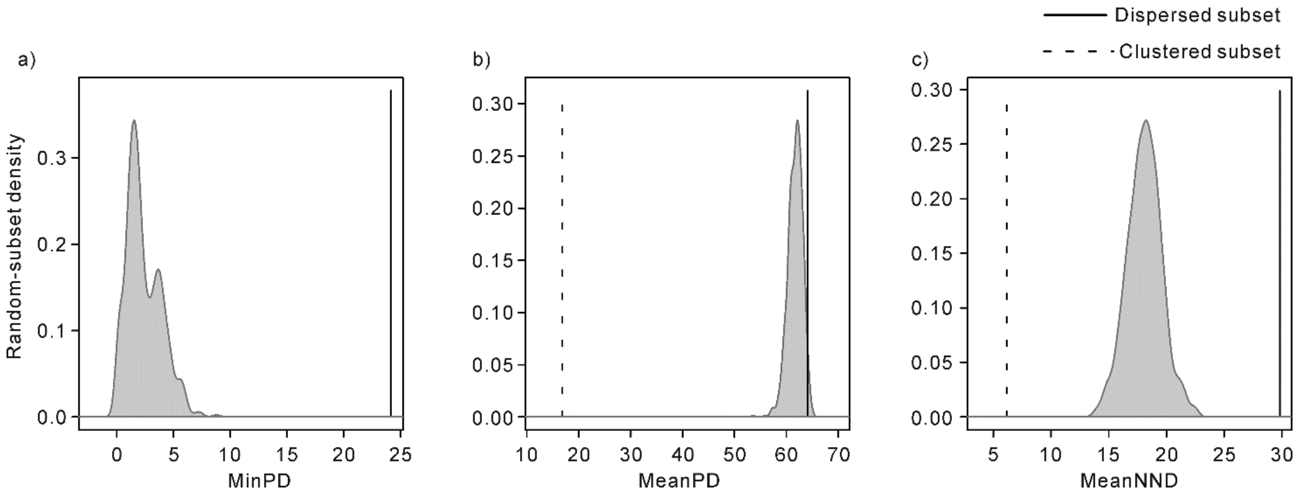

**Figure S2. Distance-based comparison of selected Cricetidae subsets with random-subset baselines.** Distance-based metrics for the selected Cricetidae subsets were compared with 1000 random subsets of the same size drawn from the same 512-species candidate pool. Panels a–c show minimum pairwise patristic distance (MinPD), mean pairwise patristic distance (MeanPD), and mean nearest-neighbor phylogenetic distance (MeanNND). For all the three metrics, larger values indicate stronger dispersion, whereas smaller values indicate stronger clustering. Vertical lines indicate observed values of the selected subsets: solid lines represent the phylogenetically dispersed subset, and dashed lines represent the clustered subset. Density curves show empirical distributions from random subsets. Maximum pairwise patristic distance (MaxPD) is not shown as a separate panel because it was constant across all random subsets in this candidate pool.

#### Section S2. Sensitivity of Subsetting to Sampling Design

The analyses included Cricetidae candidate pools of $N$ = 32, 64, 128, 256, and 512 species, and target subset sizes of $s$ = 8, 16, 32, and 64, retaining only combinations for which $s < N$. Across all feasible $N$–$s$ combinations, phylogenetically dispersed subsets showed larger MinPD, MeanPD, and MeanNND than random subsets drawn from the same candidate pool, whereas clustered subsets showed smaller MaxPD, MeanPD, and MeanNND (Table S2-S4). These contrasts were consistently significant for MinPD and MeanNND in dispersed subsets and for MaxPD, MeanPD, and MeanNND in clustered subsets. For dispersed subsets, MeanPD showed the same direction of change but was less consistently significant when the target subset represented a relatively large fraction of the candidate pool, suggesting that gains in overall mean pairwise distance were more constrained under high sampling fractions. As expected, absolute distance values changed with both $N$ and $s$, because larger candidate pools provide more opportunities to select widely separated species and larger target subsets impose stronger packing constraints. Nevertheless, despite changes in absolute distance values and some nonsignificant MeanPD comparisons for dispersed subsets, dispersed and clustered subsets remained displaced in opposite directions from the random baseline, consistent with the pattern observed in the Cricetidae analysis.

The nested design also helped separate the effects of absolute subset size from the effects of sampling fraction, as illustrated in Table S2-S4. Holding $N$ fixed and increasing $s$ generally reduced the ability of the dispersed subset to maintain large nearest-neighbor distances, because more species had to be accommodated within the same candidate pool. Holding $s$ fixed and increasing $N$ generally made stronger dispersion possible, because the selection algorithm could draw from a broader set of phylogenetic positions. In a few cases, however, nested candidate pools produced identical selected subsets across adjacent pool sizes; for example, the same dispersed 8-species subset was recovered from the $N$ = 128 and $N$ = 256 pools, resulting in identical MeanNND values.

Comparisons with the same sampling fraction but different absolute $N$ and $s$ further indicated that sampling fraction alone did not fully determine subset behavior. For example, although $N$ = 32, $s$ = 16 and $N$ = 64, $s$ = 32 both represent a sampling fraction of 0.5, the clustered subset was not significant in the former MeanNND comparison but was significant in the latter. This difference suggests that the achievable level of clustering or dispersion depends not only on $s/N$, but also on absolute subset size and the phylogenetic scale and structure of the candidate pool.

#### Supplementary Tables

##### Tables S1. Species lists and subset memberships used in the analyses.

This table is deposited in Dryad (https://doi.org/10.5061/dryad.xxxxxxx)and is not included in this Supplementary Materials file.

##### Table S2. Boundary-distance criteria in the nested Cricetidae sensitivity analysis.

| Subset size (*s*) | *N* = 32 | *N* = 64 | *N* = 128 | *N* = 256 | *N* = 512 |
|---|---|---|---|---|---|
| Panel A: MinPD for phylogenetically dispersed subsets | | | | | |
| 8 | 41.83 | 47.62 | 51.95 | 51.95 | 53.56 |
| 16 | 30.45 | 33.50 | 36.77 | 39.25 | 40.81 |
| 32 | — | 25.55 | 27.47 | 30.29 | 32.44 |
| 64 | — | — | 17.45 | 20.69 | 24.18 |
| Panel B: MaxPD for phylogenetically clustered subsets | | | | | |
| 8 | 33.50 | 17.05 | 10.90 | 5.53 | 5.65 |
| 16 | 51.95 | 33.50 | 20.25 | 11.31 | 7.98 |
| 32 | – | 41.83 | 33.50 | 20.25 | 11.31 |
| 64 | – | – | 41.83 | 33.50 | 20.25 |

Values are observed boundary-distance values. All reported values were significant relative to random subsets generated from the same candidate pool and with the same target subset size at empirical one-sided $p < 0.001$. For Panel A, significance was evaluated in the direction of larger minimum pairwise patristic distance (MinPD) values than random subsets. For Panel B, significance was evaluated in the direction of smaller maximum pairwise patristic distance (MaxPD) values than random subsets. Repeated MinPD values across some *N*-*s* combinations are expected under the nested candidate-pool design when the dispersed-subset procedure recovers subsets bounded by the same most distant pair. En dashes indicate combinations not analyzed because $s \geq N$.

**Table S3. MeanPD in the nested Cricetidae sensitivity analysis.**

| Subset type | Subset size (*s*) | *N* = 32 | *N* = 64 | *N* = 128 | *N* = 256 | *N* = 512 |
|---|---|---|---|---|---|---|
| Dispersed | 8 | 71.33** | 72.56*** | 74.02*** | 74.02*** | 74.26*** |
| Dispersed | 16 | 64.86 | 68.39*** | 68.94** | 69.33*** | 71.43*** |
| Dispersed | 32 | – | 63.35 | 63.95 | 65.14* | 67.34*** |
| Dispersed | 64 | – | – | 62.45 | 62.85 | 64.11* |
| Clustered | 8 | 22.74*** | 13.16*** | 7.17*** | 4.35*** | 4.19*** |
| Clustered | 16 | 40.09*** | 23.11*** | 16.63*** | 8.13*** | 6.15*** |
| Clustered | 32 | — | 37.51*** | 24.02*** | 15.90*** | 9.34*** |
| Clustered | 64 | — | — | 37.37*** | 23.47*** | 16.85*** |

Values are shown as observed mean pairwise patristic distance (MeanPD) values. Asterisks indicate empirical one-sided significance relative to random subsets generated from the same candidate pool and with the same target subset size: * $p < 0.05$, ** $p < 0.01$, and *** $p < 0.001$. For dispersed subsets, significance was evaluated in the direction of larger MeanPD than random subsets; for clustered subsets, significance was evaluated in the direction of smaller MeanPD than random subsets. En dashes indicate combinations not analyzed because $s \geq N$. The identical dispersed MeanPD values for $N = 128$ and $N = 256$ at $s = 8$ reflect recovery of the same selected subset from the nested candidate pools.

**Table S4. MeanNND in the nested Cricetidae sensitivity analysis.**

| Subset type | Subset size (*s*) | *N* = 32 | *N* = 64 | *N* = 128 | *N* = 256 | *N* = 512 |
|---|---|---|---|---|---|---|
| Dispersed | 8 | 54.36*** | 54.46** | 58.65*** | 58.65*** | 60.35*** |
| Dispersed | 16 | 40.67*** | 41.50*** | 47.14*** | 47.26*** | 48.25*** |
| Dispersed | 32 | – | 32.16*** | 35.52*** | 37.90*** | 39.09*** |
| Dispersed | 64 | – | – | 26.35*** | 28.42*** | 29.83*** |
| Clustered | 8 | 17.59*** | 7.79*** | 5.04*** | 2.98*** | 2.65*** |
| Clustered | 16 | 26.03 | 12.51*** | 9.02*** | 4.13*** | 3.26*** |
| Clustered | 32 | – | 17.57** | 9.91*** | 7.74*** | 3.23*** |
| Clustered | 64 | – | – | 13.91*** | 8.15*** | 6.18*** |

Values are shown as observed mean nearest-neighbor phylogenetic distance (MeanNND) values. Asterisks indicate empirical one-sided significance relative to random subsets generated from the same candidate pool and with the same target subset size: * $p < 0.05$, ** $p < 0.01$, and *** $p < 0.001$. For dispersed subsets, significance was evaluated in the direction of larger MeanNND than random subsets; for clustered subsets, significance was evaluated in the direction of smaller MeanNND than random subsets. En dashes indicate combinations not analyzed because $s \geq N$. The identical dispersed MeanNND values for $N = 128$ and $N = 256$ at $s = 8$ reflect recovery of the same selected subset from the nested candidate pools.

##### Table S5. MeanOffCor in the nested Cricetidae sensitivity analysis.

| Subset type | *s* | *N* = 32 | *N* = 64 | *N* = 128 | *N* = 256 | *N* = 512 |
|---|---|---|---|---|---|---|
| Dispersed | 8 | 0.080** | 0.065*** | 0.046*** | 0.046*** | 0.043*** |
| Dispersed | 16 | 0.164 | 0.118*** | 0.111** | 0.106*** | 0.079*** |
| Dispersed | 32 | – | 0.183 | 0.176 | 0.160* | 0.132*** |
| Dispersed | 64 | – | – | 0.195 | 0.190 | 0.173* |
| Clustered | 8 | 0.707*** | 0.830*** | 0.908*** | 0.944*** | 0.946*** |
| Clustered | 16 | 0.483*** | 0.702*** | 0.786*** | 0.895*** | 0.921*** |
| Clustered | 32 | — | 0.516*** | 0.690*** | 0.795*** | 0.880*** |
| Clustered | 64 | — | — | 0.518*** | 0.697*** | 0.783*** |

Values are shown as observed mean off-diagonal correlation (MeanOffCor) values. Asterisks indicate empirical one-sided significance relative to random subsets generated from the same candidate pool and with the same target subset size: * $p < 0.05$, ** $p < 0.01$, and *** $p < 0.001$. For dispersed subsets, significance was evaluated in the direction of lower MeanOffCor than random subsets; for clustered subsets, significance was evaluated in the direction of higher MeanOffCor than random subsets. En dashes indicate combinations not analyzed because subset size $s \geq N$. The identical dispersed MeanOffCor values for $N = 128$ and $N = 256$ at $s = 8$ reflect recovery of the same selected subset from the nested candidate pools.

##### Table S6. MaxOffCor in the nested Cricetidae sensitivity analysis.

| Subset type | *s* | *N* = 32 | *N* = 64 | *N* = 128 | *N* = 256 | *N* = 512 |
|---|---|---|---|---|---|---|
| Dispersed | 8 | 0.461*** | 0.386*** | 0.330*** | 0.330*** | 0.309*** |
| Dispersed | 16 | 0.607*** | 0.568*** | 0.526*** | 0.494*** | 0.474*** |
| Dispersed | 32 | – | 0.671*** | 0.646*** | 0.610*** | 0.582*** |
| Dispersed | 64 | – | – | 0.775*** | 0.733*** | 0.688*** |
| Clustered | 8 | 0.909 | 0.960* | 0.983** | 0.983** | 0.991** |
| Clustered | 16 | 0.866 | 0.980 | 0.960 | 0.983* | 0.983* |
| Clustered | 32 | — | 0.939 | 0.980 | 0.974 | 0.984 |
| Clustered | 64 | — | — | 0.983 | 0.980 | 0.991 |

Values are shown as observed maximum off-diagonal correlation (MaxOffCor) values. Asterisks indicate empirical one-sided significance relative to random subsets generated from the same candidate pool and with the same target subset size: * $p < 0.05$, ** $p < 0.01$, and *** $p < 0.001$. For dispersed subsets, significance was evaluated in the direction of lower MaxOffCor than random subsets; for clustered subsets, significance was evaluated in the direction of higher MaxOffCor than random subsets. For clustered subsets, MaxOffCor remains a valid summary but is less discriminating at larger subset sizes, because random subsets may already contain at least one highly correlated species pair; therefore, nonsignificant clustered-subset MaxOffCor values do not necessarily indicate weak clustering. En dashes indicate combinations not analyzed because subset size $s \geq N$. The identical dispersed MaxOffCor values for $N = 128$ and $N = 256$ at $s = 8$ reflect recovery of the same selected subset from the nested candidate pools.

**Table S7. PIESS for RMSE in the nested Cricetidae sensitivity analysis.**

| Subset type | *s* | *N* = 32 | *N* = 64 | *N* = 128 | *N* = 256 | *N* = 512 |
|---|---|---|---|---|---|---|
| Dispersed | 8 | 7.12** | 7.38*** | 7.69*** | 7.59*** | 7.70*** |
| Dispersed | 16 | 8.66** | 9.55*** | 10.35*** | 10.35*** | 12.06*** |
| Dispersed | 32 | — | 10.09* | 10.71** | 11.60*** | 13.09*** |
| Dispersed | 64 | — | — | 10.60* | 11.74*** | 12.96*** |
| Clustered | 8 | <4* | <4 | <4 | <4 | <4 |
| Clustered | 16 | 4.65*** | <4*** | <4*** | <4*** | <4*** |
| Clustered | 32 | — | 4.76*** | <4*** | <4*** | <4*** |
| Clustered | 64 | — | — | 5.13*** | <4*** | <4*** |

Values are shown as observed prediction-metric-based independence-equivalent sample size (PIESS) values for predictive RMSE. Asterisks indicate empirical one-sided significance relative to random subsets generated from the same candidate pool and with the same target subset size: * $p < 0.05$, ** $p < 0.01$, and *** $p < 0.001$. For dispersed subsets, significance was evaluated in the direction of higher PIESS than random subsets; for clustered subsets, significance was evaluated in the direction of lower PIESS than random subsets. En dashes indicate combinations not analyzed because subset size ($s$) $\geq N$. Values are rounded to two decimal places; identical displayed values may arise from rounding, e.g. underlying values of 10.3472 and 10.3518, which are distinct at higher precision, both appear as 10.35 after rounding.

**Table S8. PIESS for MAE in the nested Cricetidae sensitivity analysis.**

| Subset type | *s* | *N* = 32 | *N* = 64 | *N* = 128 | *N* = 256 | *N* = 512 |
|---|---|---|---|---|---|---|
| Dispersed | 8 | 6.92** | 7.24*** | 7.59*** | 7.37*** | 7.59*** |
| Dispersed | 16 | 7.93* | 9.25** | 10.27*** | 9.98*** | 11.63*** |
| Dispersed | 32 | — | 8.78 | 9.97** | 10.61*** | 12.31*** |
| Dispersed | 64 | — | — | 9.77 | 10.40** | 11.54*** |
| Clustered | 8 | <4 | <4 | <4 | <4 | <4 |
| Clustered | 16 | <4*** | <4** | <4** | <4** | <4** |
| Clustered | 32 | — | <4*** | <4*** | <4*** | <4*** |
| Clustered | 64 | — | — | <4*** | <4*** | <4*** |

Values are shown as observed prediction-metric-based independence-equivalent sample size (PIESS) values for predictive MAE. Asterisks indicate empirical one-sided significance relative to random subsets generated from the same candidate pool and with the same target subset size: * $p < 0.05$, ** $p < 0.01$, and *** $p < 0.001$. For dispersed subsets, significance was evaluated in the direction of higher PIESS than random subsets; for clustered subsets, significance was evaluated in the direction of lower PIESS than random subsets. En dashes indicate combinations not analyzed because subset size ($s$) $\geq N$. Minor deviations from monotonic trends across subset sizes are due to stochastic fluctuations in finite-sample estimates across simulation replicates. Values are rounded to two decimal places; identical displayed values may arise from rounding.

**Table S9. MeanOffCor under alternative covariance assumptions in the nested Cricetidae sensitivity analysis.**

| Covariance setting | $N$ = 128, $s$ = 8 | $N$ = 128, $s$ = 64 | $N$ = 512, $s$ = 8 | $N$ = 512, $s$ = 64 |
|---|---|---|---|---|
| Panel A. $\lambda$-transformed BM covariance structures | | | | |
| $\lambda$ = 0.00 | 0.000/0.000 | 0.000/0.000 | 0.000/0.000 | 0.000/0.000 |
| $\lambda$ = 0.25 | 0.011***/0.227*** | 0.049/0.130*** | 0.011***/0.237*** | 0.043*/0.196*** |
| $\lambda$ = 0.50 | 0.023***/0.454*** | 0.097/0.259*** | 0.021***/0.473*** | 0.087*/0.391*** |
| $\lambda$ = 0.75 | 0.034***/0.681*** | 0.146/0.389*** | 0.032***/0.710*** | 0.130*/0.587*** |
| $\lambda$ = 1.00 | 0.046***/0.908*** | 0.195/0.518*** | 0.043***/0.946*** | 0.173*/0.783*** |
| Panel B. OU covariance structures | | | | |
| $h/H$ = 0.05 | <0.001***/0.113*** | <0.001***/0.002*** | <0.001***/0.252*** | <0.001***/0.012*** |
| $h/H$ = 0.10 | <0.001***/0.306*** | 0.001***/0.010*** | <0.001***/0.486*** | <0.001***/0.069*** |
| h/H = 0.25 | 0.002***/0.607*** | 0.027***/0.084*** | 0.002***/0.744*** | 0.020***/0.311*** |
| h/H = 0.50 | 0.012***/0.762*** | 0.081*/0.228*** | 0.011***/0.853*** | 0.067***/0.524*** |
| h/H = 1.00 | 0.025***/0.841*** | 0.131/0.358*** | 0.023***/0.904*** | 0.113**/0.656*** |
| Panel C. EB covariance structures | | | | |
| $\rho = -4.0$ | 0.125*** / 0.991*** | 0.353 / 0.882*** | 0.121*** / 0.995*** | 0.333 / 0.973*** |
| $\rho = -2.0$ | 0.085*** / 0.968*** | 0.286 / 0.739*** | 0.081*** / 0.982*** | 0.264 / 0.914*** |
| $\rho = -1.0$ | 0.065*** / 0.943*** | 0.243 / 0.635*** | 0.061*** / 0.968*** | 0.220 / 0.858*** |
| $\rho = -0.5$ | 0.055*** / 0.927*** | 0.219 / 0.578*** | 0.051*** / 0.958*** | 0.197* / 0.823*** |
| $\rho$ = 0.0 | 0.046*** / 0.908*** | 0.195 / 0.518*** | 0.043*** / 0.946*** | 0.173* / 0.783*** |

Each cell reports mean off-diagonal correlation (MeanOffCor) for the phylogenetically dispersed subset and the phylogenetically clustered subset, respectively, in the format dispersed/clustered. Values smaller than 0.001 are reported as < 0.001, except for the $\lambda$ = 0 row in Panel A, where MeanOffCor is theoretically zero under the independence benchmark. Asterisks indicate empirical one-sided significance relative to random subsets generated from the same candidate pool and with the same target subset size: * $p$ < 0.05, ** $p$ < 0.01, and *** $p$ < 0.001. For dispersed subsets, significance was evaluated in the direction of lower MeanOffCor than random subsets; for clustered subsets, significance was evaluated in the direction of higher MeanOffCor than random subsets. Parameter definitions and interpretations are provided in Supplementary Methods Section S4.

**Table S10. MaxOffCor under alternative covariance assumptions in the nested Cricetidae sensitivity analysis.**

| Covariance setting | $N$ = 128, $s$ = 8 | $N$ = 128, $s$ = 64 | $N$ = 512, $s$ = 8 | $N$ = 512, $s$ = 64 |
|---|---|---|---|---|
| Panel A. $\lambda$-transformed BM covariance structures | | | | |
| $\lambda$ = 0.00 | 0.000/0.000 | 0.000/0.000 | 0.000/0.000 | 0.000/0.000 |
| $\lambda$ = 0.25 | 0.083***/0.246** | 0.194***/0.246 | 0.077***/0.248** | 0.172***/0.248 |
| $\lambda$ = 0.50 | 0.165***/0.492** | 0.388***/0.492 | 0.155***/0.495** | 0.344***/0.495 |
| $\lambda$ = 0.75 | 0.248***/0.737** | 0.581***/0.737 | 0.232***/0.743** | 0.516***/0.743 |
| $\lambda$ = 1.00 | 0.330***/0.983** | 0.775***/0.983 | 0.309***/0.991** | 0.688***/0.991 |
| Panel B. OU covariance structures | | | | |
| $h/H$ = 0.05 | <0.001***/0.627** | 0.002***/0.627 | <0.001***/0.774** | <0.001***/0.774 |
| $h/H$ = 0.10 | <0.001***/0.792** | 0.044***/0.792 | <0.001***/0.880** | 0.013***/0.880 |
| $h/H$ = 0.25 | 0.021***/0.911** | 0.285***/0.911 | 0.018***/0.950** | 0.174***/0.950 |
| $h/H$ = 0.50 | 0.100***/0.951** | 0.505***/0.951 | 0.091***/0.973** | 0.383***/0.973 |
| $h/H$ = 1.00 | 0.194***/0.969** | 0.643***/0.969 | 0.179***/0.983** | 0.532***/0.983 |
| Panel C. EB covariance structures | | | | |
| $\rho = -4.0$ | 0.747*** / 0.999** | 0.973*** / 0.999 | 0.723*** / 0.999** | 0.954*** / 0.999 |
| $\rho = -2.0$ | 0.559*** / 0.995** | 0.911*** / 0.995 | 0.534*** / 0.997** | 0.865*** / 0.997 |
| $\rho = -1.0$ | 0.445*** / 0.990** | 0.853*** / 0.990 | 0.421*** / 0.995** | 0.787*** / 0.995 |
| $\rho = -0.5$ | 0.387*** / 0.987** | 0.817*** / 0.987 | 0.364*** / 0.993** | 0.740*** / 0.993 |
| $\rho = 0.0$ | 0.330*** / 0.983** | 0.775*** / 0.983 | 0.309*** / 0.991** | 0.688*** / 0.991 |

Each cell reports maximum off-diagonal correlation (MaxOffCor) values for the phylogenetically dispersed subset and the phylogenetically clustered subset, respectively, in the format dispersed / clustered. Values smaller than 0.001 are reported as < 0.001, except for the $\lambda$ = 0 row in Panel A, where MaxOffCor is theoretically zero under the independence benchmark. Asterisks indicate empirical one-sided significance relative to random subsets generated from the same candidate pool and with the same target subset size: * $p$ < 0.05, ** $p$ < 0.01, and *** $p$ < 0.001. For dispersed subsets, significance was evaluated in the direction of lower MaxOffCor than random subsets; for clustered subsets, significance was evaluated in the direction of higher MaxOffCor than random subsets. MaxOffCor is most informative for dispersed subsets, because it evaluates whether the selected subset still contains a strongly correlated species pair. For clustered subsets, it remains a valid summary but is less discriminating at larger subset sizes, because random subsets may already contain at least one highly correlated species pair; therefore, nonsignificant clustered-subset MaxOffCor values for $s$ = 64 do not necessarily indicate weak clustering. If clustered subsets were the primary object of analysis, minimum off-diagonal correlation would provide a more targeted diagnostic, because it evaluates whether even the least correlated pair within a subset remains relatively dependent. Parameter definitions and interpretations are provided in Supplementary Methods Section S4.

**Table S11. PIESS for RMSE under alternative covariance assumptions in the nested Cricetidae sensitivity analysis.**

| Covariance setting | N = 128, s = 8 | N = 128, s = 64 | N = 512, s = 8 | N = 512, s = 64 |
|---|---|---|---|---|
| Panel A. $\lambda$-transformed BM covariance structures | | | | |
| $\lambda$ = 0.00 | 8.00 / 8.20 | 62.81 / 62.25 | 8.07 / 8.15 | 62.07 / 62.46 |
| $\lambda$ = 0.25 | 8.05* / 6.45*** | 47.72* / 31.88*** | 7.71 / 6.46*** | 50.67*** / 21.87*** |
| $\lambda$ = 0.50 | 7.81** / <4*** | 27.11 / 14.17*** | 8.03*** / <4*** | 29.89** / 8.35*** |
| $\lambda$ = 0.75 | 7.75*** / <4** | 16.13* / 8.06*** | 7.92*** / <4** | 19.86*** / 4.06*** |
| $\lambda$ = 1.00 | 7.45*** / <4 | 10.81** / 4.67*** | 7.73*** / <4 | 12.17*** / <4*** |
| Panel B. OU covariance structures | | | | |
| $h/H$ = 0.05 | 8.01 / 7.15*** | 62.59 / 62.26 | 7.90 / 5.79*** | 62.14 / 57.00* |
| $h/H$ = 0.10 | 8.00 / 4.99*** | 62.40* / 54.29** | 7.98 / <4*** | 62.98*** / 36.38** |
| $h/H$ = 0.25 | 8.01 / <4*** | 53.22*** / 32.72*** | 8.12* / <4*** | 57.97*** / 10.08*** |
| $h/H$ = 0.50 | 8.05*** / <4*** | 28.59*** / 15.17*** | 7.90** / <4*** | 35.51*** / 4.74*** |
| $h/H$ = 1.00 | 7.85*** / <4** | 19.01*** / 8.40*** | 7.78** / <4** | 21.54*** / <4*** |
| Panel C. EB covariance structures | | | | |
| $\rho = -4.0$ | 5.84*** / <4 | <4 / <4 | 5.96*** / <4 | 5.84*** / <4 |
| $\rho = -2.0$ | 7.10*** / <4 | 5.76 / <4*** | 7.14*** / <4 | 7.10*** / <4 |
| $\rho = -1.0$ | 7.41*** / <4 | 7.65 / <4*** | 7.52*** / <4 | 7.41*** / <4 |
| $\rho = -0.5$ | 7.71*** / <4 | 8.99** / <4*** | 7.49*** / <4 | 7.71*** / <4 |
| $\rho = 0.0$ | 7.69*** / <4 | 10.81** / 5.13*** | 7.73*** / <4 | 7.69*** / <4 |

Each cell reports prediction-metric-based independence-equivalent sample size (PIESS) values for RMSE for the phylogenetically dispersed subset and the phylogenetically clustered subset, respectively, in the format dispersed / clustered. Asterisks indicate empirical one-sided significance relative to random subsets generated from the same candidate pool and with the same target subset size: * $p < 0.05$, ** $p < 0.01$, and *** $p < 0.001$. For dispersed subsets, significance was evaluated in the direction of higher PIESS than random subsets; for clustered subsets, significance was evaluated in the direction of lower PIESS than random subsets. In Panel A, $\lambda = 0$ represents the limiting case in which all off-diagonal phylogenetic covariance is removed; under this condition, PIESS is expected to be close to the nominal subset size, with small deviations possible because PIESS is estimated from finite simulation replicates and interpolation against an empirical benchmark curve. Parameter definitions and interpretations are provided in Supplementary Methods Section S4.

**Table S12. PIESS for MAE under alternative covariance assumptions in the nested Cricetidae sensitivity analysis.**

| Covariance setting | N = 128, s = 8 | N = 128, s = 64 | N = 512, s = 8 | N = 512, s = 64 |
|---|---|---|---|---|
| Panel A. $\lambda$-transformed BM covariance structures | | | | |
| $\lambda$ = 0.00 | 7.74 / 8.09 | 67.41* / 62.37 | 7.82 / 7.96 | 61.15 / 62.44 |
| $\lambda$ = 0.25 | 7.88* / 5.84*** | 45.48 / 31.94*** | 7.61 / 5.90*** | 51.53*** / 19.88*** |
| $\lambda$ = 0.50 | 7.73** / <4*** | 25.66 / 12.49*** | 7.78*** / <4*** | 28.73** / 6.92*** |
| $\lambda$ = 0.75 | 7.69*** / <4* | 15.54 / 6.55*** | 7.85*** / <4* | 17.76*** / <4*** |
| $\lambda$ = 1.00 | 7.38*** / <4 | 10.14* / <4*** | 7.59*** / <4 | 10.62** / <4*** |
| Panel B. OU covariance structures | | | | |
| $h/H$ = 0.05 | 7.91 / 7.04*** | 67.06 / 62.61 | 7.71 / 5.48*** | 61.39 / 56.06 |
| $h/H$ = 0.10 | 7.89 / 4.43*** | 62.13 / 55.40* | 7.86 / <4*** | 67.14** / 37.60*** |
| $h/H$ = 0.25 | 7.88 / <4*** | 53.99*** / 32.33*** | 7.89 / <4*** | 58.58*** / 8.41*** |
| $h/H$ = 0.50 | 7.87*** / <4*** | 29.67*** / 13.97*** | 7.83** / <4*** | 35.29*** / <4*** |
| $h/H$ = 1.00 | 7.73*** / <4* | 17.17*** / 7.28*** | 7.59** / <4* | 19.90*** / <4*** |
| Panel C. EB covariance structures | | | | |
| $\rho = -4.0$ | 5.77*** / <4 | <4 / <4 | 5.88*** / <4 | <4 / <4 |
| $\rho = -2.0$ | 6.77*** / <4 | 5.29 / <4** | 6.88*** / <4 | 5.84* / <4* |
| $\rho = -1.0$ | 7.28*** / <4 | 6.91 / <4*** | 7.30*** / <4 | 7.76* / <4*** |
| $\rho = -0.5$ | 7.42*** / <4 | 7.99 / <4*** | 7.36*** / <4 | 9.69*** / <4*** |
| $\rho = 0.0$ | 7.51*** / <4 | 10.16* / <4*** | 7.58*** / <4 | 11.18*** / <4*** |

Each cell reports prediction-metric-based independence-equivalent sample size (PIESS) values for MAE for the phylogenetically dispersed subset and the phylogenetically clustered subset, respectively, in the format dispersed / clustered. Asterisks indicate empirical one-sided significance relative to random subsets generated from the same candidate pool and with the same target subset size: * $p < 0.05$, ** $p < 0.01$, and *** $p < 0.001$. For dispersed subsets, significance was evaluated in the direction of higher PIESS than random subsets; for clustered subsets, significance was evaluated in the direction of lower PIESS than random subsets. In Panel A, $\lambda = 0$ represents the limiting case in which all off-diagonal phylogenetic covariance is removed; under this condition, PIESS is expected to be close to the nominal subset size, with small deviations possible because PIESS is estimated from finite simulation replicates and interpolation against an empirical benchmark curve. Parameter definitions and interpretations are provided in Supplementary Methods Section S4.
